# Missing links connect the phylogeographic structure of endangered red pandas, remaining as one species – *Ailurus fulgens*, and expediting conservation

**DOI:** 10.1101/2024.08.27.609966

**Authors:** Lucy A. Dueck, Deniz Aygören Uluer

## Abstract

Monitoring biodiversity depends on well-informed taxonomy, reflecting the underlying biology of organisms critical for appropriate conservation management. The taxonomy of red pandas, endangered mammals distributed along a ∼2500km montane range fringing the southern Tibetan Plateau, has been perplexing. Finally resolved as evolutionarily unique Musteloidea, further division of *Ailurus* into two geographic variants remains contentious. Red pandas are threatened by extinction from anthropogenic disturbances with consequent population decline; continued gene flow is therefore crucial to maintain adaptive potential. A recent phylogenomic study from sampling range ends and a constrictive species concept split red pandas into two species. Subsequent studies supplied additional data missing from their midrange. We evaluated GenBank mtDNA sequences from 393 animals, plotted sampling locations, and analyzed with and without midrange samples. Two sublineages of monophyletic *Ailurus* were weakly supported by one method when the midrange was excluded, but not when it was included. Using more stringent analyses, *Ailurus* was strongly confirmed as a single species in all circumstances, although the western sublineage (*A. f. fulgens*) was weakly supported within. Four haplogroups associated with specific regions, suggesting broad phylogeographic patterning and gene flow, with sympatric crossover and a cline bracketing unsampled Bhutan in the midrange. Red pandas are due for IUCN Red List reassessment in 2025; unwarranted taxonomic revision would delay and imperil action needed to prevent extinction. We recommend maintaining *Ailurus* as one species, recognizing and protecting biodiversity at one intraspecific level, allowing for gene flow in conservation management, and extensively investigating intergradation of midrange red pandas.

## Introduction

Conservation biology and taxonomy are inextricably linked in the struggle to understand and preserve biodiversity during the current global extinction crisis. Development of action strategies for conservation depends on accurate, complete information about groups of organisms (Dubois 2003; Ely et al. 2017). While taxonomy has benefitted from expansion of morphological to genetic data (McNeely 2002), both forms are complementary to taxonomic conclusions, also reliant on the concept and definition of “species” applied (Frankham et al. 2012). The correct classification of a group of organisms informs on its basic biology – specific knowledge about that group essential to guide appropriate conservation policy and ensure the group’s persistence when management becomes necessary but resources are limited. The depth of differences between taxonomic groups is used to decide many management strategies (Frankham et al. 2012). In turn, having correct and complete biological information is essential to inform on taxonomic delimitation, and especially so for organisms of conservation concern. Such information is acquired via good scientific methodology by thoroughly sampling many individuals throughout the native range, especially at prospective contact zones (Chambers et al. 2023). Knowledge of ecology, morphology, behavior, reproduction, gene flow potential, geographic variation, distribution, as well as genetics, are key factors in determining distinctiveness between closely related groups.

Case in point is the endangered red panda, *Ailurus fulgens*, a smaller arctoid carnivore that evolved to feed primarily on bamboo. Fossil records indicate ancestral linkages throughout Eurasia and North America in a Holarctic distribution (Roberts and Gittleman 1984; Salesa et al. 2011) dated back 25-35mya via molecular data (Nyakatura and Bininda-Emonds 2012). But today, red pandas are the sole representative of family Ailuridae, thought to have diverged 2-3mya (Li et al. 2005; Wallace 2011; Hu et al. 2020), part of a trichotomy within a broad Musteloidea clade (Flynn et al. 2000; Arnason et al. 2007; Sato et al. 2009). The taxonomic uniqueness of an extant ailurid thus makes them evolutionarily invaluable and especially worthy of conservation (Redding and Mooers 2006). The species *A. fulgens* was first described by Cuvier in 1825 from Nepal at the western end of its range, then again by Thomas in 1902 from China at the eastern end of its range. It was therefore split into different subspecies (*A. f. fulgens* and *A. f. styani*, respectively) based on this separated distribution and geographical variation in morphology and pelage (Roberts 1982). Red pandas are now known to be distributed in semi-isolated populations along a ragged, continuous arc spanning ∼2500km, narrow in the west (125km) but widening to 700km in eastern regions, at the southern edge of the Qinghai-Tibetan Plateau in the Himalayan foothills of Nepal, Tibet, India, Bhutan, and Myanmar, and in the Hengduan and other mountain ranges of Yunnan and Sichuan, China. They inhabit a specialized ecological niche within mixed deciduous-coniferous forests having a bamboo understory, in temporate montane regions at elevations of ∼1500-4800m, variable by region. They subsist on a diet of bamboo leaves and shoots, supplemented by occasional berries, bird eggs, and insects. They are arboreal, elusive, and generally solitary except during breeding season (Yonzon and Hunter 1991; Choudhury 2001; Pradham et al. 2001; Dorji et al. 2012; Glatston et al. 2015; Thapa et al. 2018; Dendup et al. 2023). They are threatened with extinction due mainly to anthropogenic disturbances such as habitat destruction and fragmentation, farming, grazing, poaching, and dogs (Yonzon and Hunter 1991; Wei et al. 1999a; Choudhury 2001; Thapa et al. 2018). Red pandas are therefore listed as Endangered A2 from rapid population decline (≥ 50% in last three generations) by the IUCN Red List, version 2023-1 (Glatston et al. 2015), which identifies taxa needing targeted efforts to prevent extinction.

Official attempts at understanding their intraspecific variability began with Pocock’s (1941) compendium of morphological differences between subspecies, but he found too much variation within to justify full species status. Roberts (1982) continued morphological investigations – although he observed some differences in 6-7 characters, the overall coefficient of difference was not high enough to validate even subspecies status. Gentz (1989) initiated genetic analysis with an allozyme study from both groups – 23 of 25 loci were monomorphic so calculated indices of distance and similarity were outside separate species but within subspecies range. One DNA fingerprinting study (Wei et al. 1999b) in China included samples from both subspecies and two hybrid crosses; they found one heritable band characteristic for each subspecies which were also shared by the hybrid offspring. However, all of these studies were based on a limited/unequal number of samples from both subspecies and/or collection locations. The first published mitochondrial DNA (mtDNA) study on red pandas was also conducted under these limiting circumstances (*N* = 53, all from Sichuan and Yunnan) and not surprisingly showed little variation and phylogeographic structure (Su et al. 2001).

Meanwhile, American zoos which managed captive red pandas under the Species Survival Plan® (SSP) had separated the subspecies (obtained from both ends of native red panda range) but became concerned about inbreeding within each of their small subpopulations. A mtDNA survey was therefore conducted to see if the two subpopulations were separate taxonomic units or could be interbred. Results were presented as a poster in 2001 (Dueck et al.) and informal talk to the SSP in 2002, but not published until 2021 (Dueck). This study determined that the two SSP subpopulations differed genetically somewhat, but that thorough sampling throughout the native range was necessary to draw firm taxonomic conclusions about the entire species, and it was useful to establish standards comparing groups from range ends.

During that interval, two mtDNA studies were published on red pandas that expanded sampling from Sichuan/Yunnan to Tibet and Myanmar (Li et al. 2005; Hu et al. 2011); these also did not show definitive patterns of differentiation between subspecies. Two subsequent studies continued to extend sampling westward. Dalui et al. (2021) enlarged the sampling area to the midrange in India bracketing Bhutan; they found that samples from far northeast Arunachal Pradesh aligned with *A. f. styani*, while the remainder from either side of Bhutan formed a separate clade, although no support values were provided. Dueck and Steffens (2022) published their study from sampling in Nepal during 2003 – not only were regional differences among groups of *A. f. fulgens* found, but also evidence of *A. f. styani* gene flow into eastern Nepal.

All these studies were based upon sequencing 236-553 bp of mtDNA “D-loop” or control region (CR), the most variable section of that molecule. Mitochondrial DNA is useful in studies on evolutionary biology and population structure because: 1) its reduced effective population size and high mutation rate show subdivision more rapidly than nuclear DNA (Birky et al. 1989; Avise 2004), 2) gene trees from mtDNA may more accurately reflect a species tree (Moore 1995), 3) hypervariability in non-coding regions means less DNA is needed for diagnosis, and 4) since there are higher copy numbers per cell, it is more likely to amplify under difficult conditions (Taberlet et al. 1999), a valuable benefit to non-invasive sampling of endangered species. In fact, a recent study of Asian elephants, another endangered species with small fragmented populations, found mtDNA-CR preferable for determining population distinctiveness because of direct comparability, over microsatellites due to their difficult interpretation (Budd et al. 2023).

Concurrently with the later mtDNA-CR studies, two other publications, both of which invoked the Phylogenetic Species Concept that does not recognize subspecies (Nixon and Wheeler 1990; Haig et al. 2006), declared red pandas to be two separate species. Groves (2011) based his conclusion on morphological characters of skull measurements from 13-20 specimens and some photographs of pelage differences. Hu et al. (2020) conducted a study of red pandas captured from the two ends of their native range (Nepal+Tibet, Myanmar+Sichuan/Yunnan) based on genomic analyses of three marker systems – nuclear SNPs, Y-chromosome SNPs, and whole mtDNA (excluding the CR). They elevated the taxonomic rank of the two red panda subspecies to full species, backed by Groves’ morphological distinction. Yet Hu et al. (2020) also did not sample across the entire range, omitting the supposed contact zone of potential hybridization between the two (sub)species, i.e., India and Bhutan, deferring instead to a river as a sharp boundary separating them.

Our investigation aims to narrow that gap in sampling of red pandas from their native range by combining mtDNA-CR data from all previous studies to determine as much as is collectively known about the broad phylogeographic pattern for the entire species. This marker is the only consistent tool used since genetic analysis began in earnest. We address the hypothesis put forward by Roberts (1982) that *Ailurus fulgens* varies genetically and phenotypically along a cline, grading from one end of their geographic distribution to another.

## Methods

Comparable haplotype (*h*) sequences from all authors who have analyzed mtDNA-CR from red pandas in, or traced back to, their native range were collected via GenBank. These include: Su et al. (2001), Li et al. (2005), Hu et al. (2011), Dalui et al. (2021), and Dueck and Steffens (2022; WRP). Also included were sequences from captive SSP animals (Dueck 2021) since some of those haplotypes were likewise found in wild Nepal animals and provide multiple standards for putative subspecies from range ends, and two NCBI reference samples that additionally represent the two subspecies, *A. fulgens* [*fulgens*] (Arnason et al. 2007) and *A. f. styani* (Yonezawa et al. 2007). **Table A, Online Resource I,** summarizes the information available from all red panda studies used.

Haplotype duplicates among individual studies were identified by comparing each with a base dataset (SSP, WRP, two references) in unrooted phylogenetic analyses using the distance-based neighbor-joining method (NJ; Saitou and Nei 1987) with 10,000 bootstrap replicates, pairwise deletion, and the Tamura-Nei model to infer evolutionary distances, in MEGA4.0.2 (Tamura et al. 2007). Duplicates found in the resultant phylograms were visually confirmed as having identical bases at parsimony-informative positions in MEGA’s Data Explorer. When this combined dataset was aligned in MEGA’s ClustalW option, it was trimmed to a common 340bp of overlapping sequences. The shorter sequences of Su et al. (2001) were omitted because truncating to their length would lose too much phylogenetic information for all the others. **Table B, Online Resource I,** lists the 75 haplotypes with the parsimony-informative positions that differentiate them.

Up to ten musteloid representatives of the nearest families to Ailuridae were included as outgroups in subsequent analyses to root and balance the phylogeny. Exploratory NJ analyses were first conducted with the combined dataset of red panda haplotypes plus two (minimum necessary to provide statistical support for ingroup clustering) or 10 outgroups, to test the effect of excluding or including samples from the red panda midrange. Then one representative from each of three Musteloidea families were used as outgroups to root the final analyses.

Probability-based phylogenetic analyses were computed in XSEDE via the CIPRES Science Gateway (Miller et al. 2010; https://www.phylo.org), accessed August 15, 2023, using both maximum likelihood (ML; Felsenstein 1981) and Bayesian (Markov chain Monte-Carlo [MCMC]) inference (BI; Yang and Rannala 1997) methods. The best-fit model of nucleotide substitution (HKY+I+G) to be applied in both analyses was first determined in jModelTest 2.0 (Guindon and Gascuel 2003; Darriba et al. 2012) based on the Akaike Information Criterion (Akaike 1974). Programs employed were: RAxML 8.2.12 (Stamatakis 2014) with the “let RAxML halt bootstrapping automatically” option, the GTRGAMMA model, plus other default ML search options; and MrBayes 3.2.7a (Ronquist and Huelsenbeck 2003), with 10 million generations, sampling every 100^th^ generation with four parallel Monte Carlo Markov chain (MCMC) runs (one cold and three heated). The first 25,000 trees (25% burn-in) were discarded, while the remaining 75,000 trees used to visualize the maximum credibility tree. This was repeated twice and the convergence ensured.

A map of south-central Asia was drawn, showing the presumed native range of red pandas about the time of the early studies. Sampling locations from the four later native studies were plotted as accurately as possible on the map^1^. The Interactive Tree of Life software (Letunic and Bork 2021) was employed to produce the ML phylogenetic tree image, which was combined with this map into one figure with the subclades/groups and sampling locations color-coded to match.

A haplotype network was calculated in Network 10.2.0.0 (https://fluxus-technology.com) using the median-joining algorithm of Bandelt et al. (1999). The 75 red panda sequences were first pre-processed using the star contraction option, which yielded 44 taxa from which a network was drawn. The subclades/groups were color-coded as above.

Statistical tests to assess population genetic variation were computed in DnaSP 6.12.03 (Rozas et al. 2017), and pie diagrams for illustration prepared in Microsoft Excel. The data input file (**Table C, Online Resource I**) was slightly reduced to include native-only animals and haplotypes (*N* = 271, *h* = 71). Based on phylogenetic differentiation and/or geographic separation, four mitochondrial haplotype groups (“haplogroups”) matching the color-coding of the ML tree subclades/groups were compared. Several diversity (*Hd*, *π, S, k, ϴ_w_*) and divergence (*F_ST_* and *χ^2^* P-values) measures were calculated; population size changes and stability were assessed via tests of mutational neutrality (Tajima’s *D*, Fu’s *F_s_*) and by plotting mismatch distributions (inferred by pairwise nucleotide site differences) against expected values. The MEGA Distance module using a Kimura 2-parameter model of evolution, pairwise deletion, and uniform rates was employed to calculate pairwise sequence divergence values among haplotypes. Analysis of molecular variance (AMOVA) conducted in Arlequin ver. 3.5.2.2 (Excoffier and Lischer 2010) was used to separate variation attributable to within or between putative subspecies/among haplogroups.

## Results

Overall, for the six studies and two reference samples combined, 126 mtDNA-CR red panda haplotypes found among 393 samples were evaluated. After eliminating Su et al.’s (2001) shorter sequences, merging duplicates, and trimming, 340bp of 75 taxa from 340 samples were used in phylogenetic analyses. Of those positions, 56 sites were variable, 21 were singletons, and 35 were parsimony-informative; there was one insertion for Li36 at position 26. The segment of mtDNA used comprises 28.0% (340/1216bp) of the D-loop/CR and 2.1% (340/16495bp) of the entire mitochondrial genome.

Initial assessment for duplicate haplotypes revealed some noteworthy preliminary findings in individual studies summarized here but detailed in **Online Resource II**. The three early Chinese studies were found to contain only one haplotype each that affiliated with *A. f. fulgens* – Su21 from northwest Yunnan (**Figs. A, Ba**), Li-29 from south-central Tibet, and Hu-R28 from southeast Tibet/Arunachal Pradesh, all in *A. f. styani* territory. Dalui et al.’s (2021) subclade 1c consisting of three haplotypes, which they determined as belonging to *A. f. fulgens*, grouped here with *A. f. styani*, with one haplotype matching Dueck and Steffens’ (2022) WRP-C (**Fig. B**).

Exploratory NJ analyses examining the effect of exclusion *vs* inclusion of 15 midrange haplotypes (India - Dalui et al. 2021; WRP-C - Dueck and Steffens 2022) clearly demonstrates that red pandas are phylogenetically just one species, whether the midrange is included or not (**Fig. 1**). When only haplotypes from the range ends (*h* = 60) were used in the top trees (**Fig. 1a**), a well-supported (99%) monophyletic clade is weakly divided into two subspecies/sublineages. Support in the circular cladogram with two outgroups on the **left** side was 57% for *A. f. fulgens* and 60% for *A. f. styani*; in the radial phylogram with 10 outgroups on the **right**, support was slightly stronger (61% and 66%, respectively), with a slight gap visible between sublineages. When haplotypes from samples in the midrange were included (*h* = 75) in the bottom trees (**Fig. 1b**), there is 99-100% support for grouping of all red panda haplotypes together, with no support for subdivision within the species. In the circular cladogram (**Fig. 1b-left**), the full species clade is comprised of many small subclades, some being geographically specific (i.e., pink and orange subclades from midrange regions). No gap between subspecies is evident in the radial phylogram (**Fig. 1b-right**). There was consistently strong support (95-99%) for several inter- and intra-generic outgroup pairs in the top and bottom radial phylograms on the right.

**Fig. 1a.**
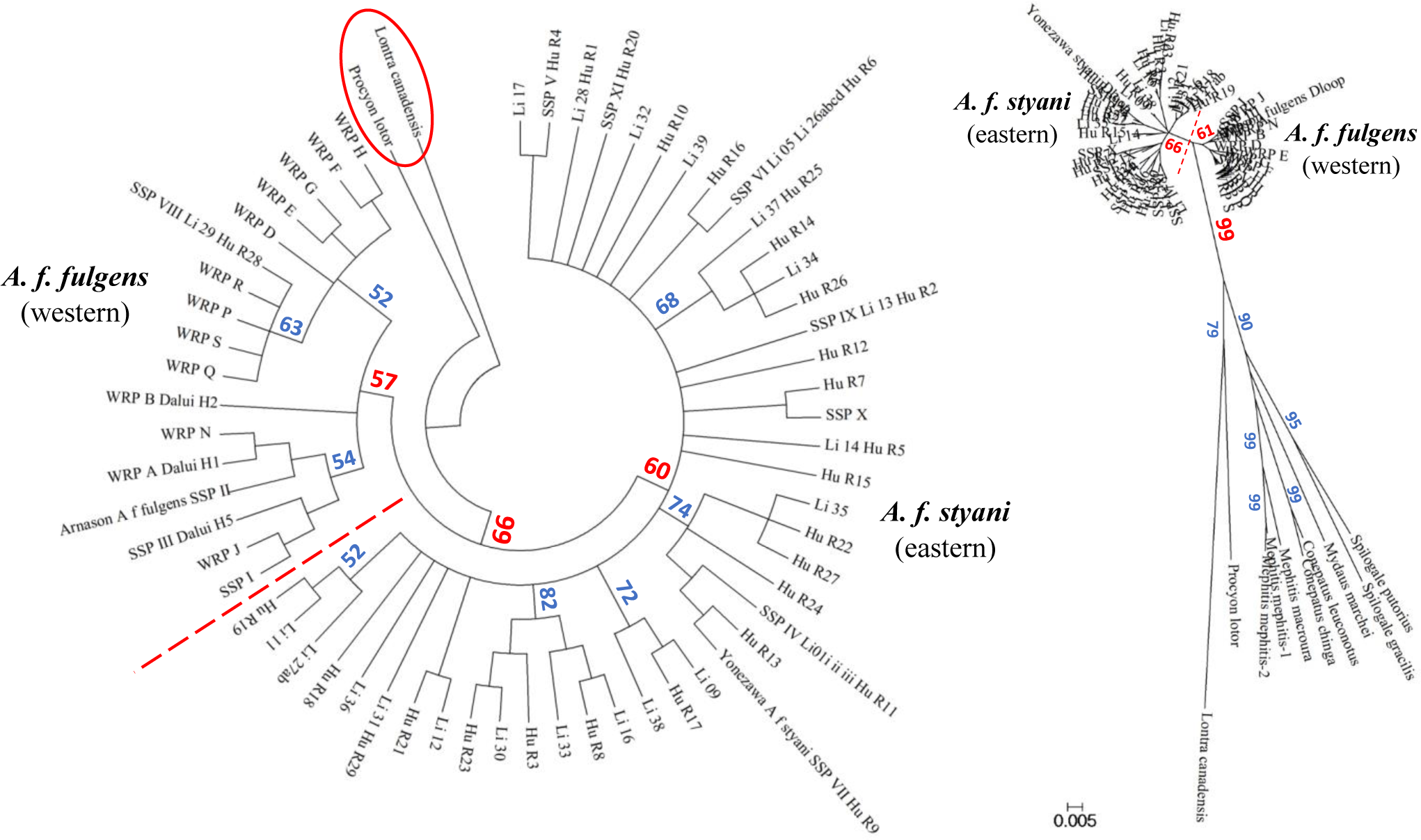
Neighbor-Joining (NJ) analysis results for all red panda mtDNA-CR haplotypes (*h* = 60) <u>excluding</u> those from the midrange (all uncombined haplotypes from India and WRP-C/Dal-H18); **-left** circular cladogram, rooted by two outgroups, depicting evolutionary relationships among 62 taxa, inferred from 10,000 bootstrap replicates in MEGA4.0.2, using pairwise deletion option and computed with the Tamura-Nei method; branches with less than 50% bootstrap support collapsed; 346 positions used in this dataset; **-right** radial phylogram, rooted by 10 outgroups, depicting evolutionary relationships among 70 taxa, computed as above using 354 positions but with no branches collapsed; tree drawn to scale, with branch lengths in the same units as the evolutionary distances to infer it (base substitutions per site)

**Fig. 1b.**
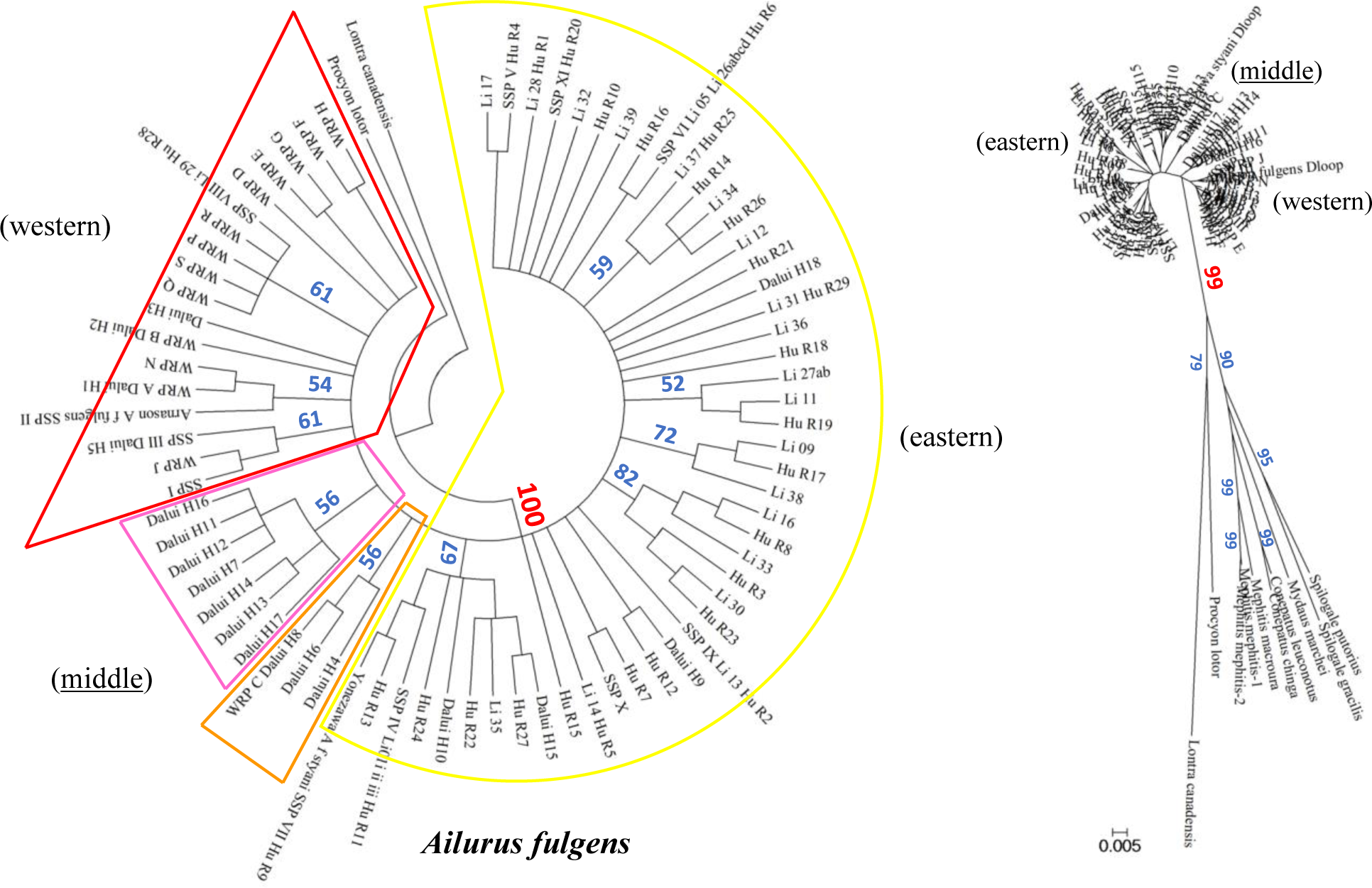
NJ analysis results for all red panda mtDNA-CR haplotypes (*h* = 75) <u>including</u> those from the midrange (Nepal, India, Myanmar, China); **-left** circular cladogram, rooted by 2 outgroups, depicting evolutionary relationships among 77 taxa, computed as above; regions of origin are highlighted by colors (red = western, pink and orange = midrange, yellow = eastern); **-right** radial phylogram, rooted by 10 outgroups, depicting evolutionary relationships among 85 taxa, computed as above

Phylogenetic analysis using the ML method yielded a tree with 92% bootstrap support for all red pandas grouping together as a monophyletic clade (**Fig. 2-left**). There was little structure within other than weak support (57%) for a larger *A. f. fulgens* subclade which was broken into two haplotype groups (“haplogroups”) also weakly supported; an unorganized group of *A. f. styani* small subclades and unpaired haplotypes was also present. The first *A. f. fulgens* haplogroup (58% support, “*Aff*-red”) contained all haplotypes previously deemed to be that subspecies (SSP - US zoos, WRP - Nepal, Arnason reference), as well as Dalui et al.’s (2021) four subclade 1b haplotypes and Li-29+Hu-R28 (Dalui-H19). On the associated map in **Fig. 2-right**, all animals from *Aff*-red were sourced to the western end of the range – Nepal, south-central Tibet, and India’s Sikkim-West Bengal states west of Bhutan – except one (Hu-R28), found in Tibet/Arunachal Pradesh east of the Siang River. But the second *A. f. fulgens* haplogroup (50% support, “*Aff*-pink”), consisting only of Dalui et al.’s (2021) seven subclade 1a haplotypes, were all from red pandas sourced to western Arunachal Pradesh, east of Bhutan but west of the Siang River.

**Fig. 2.**
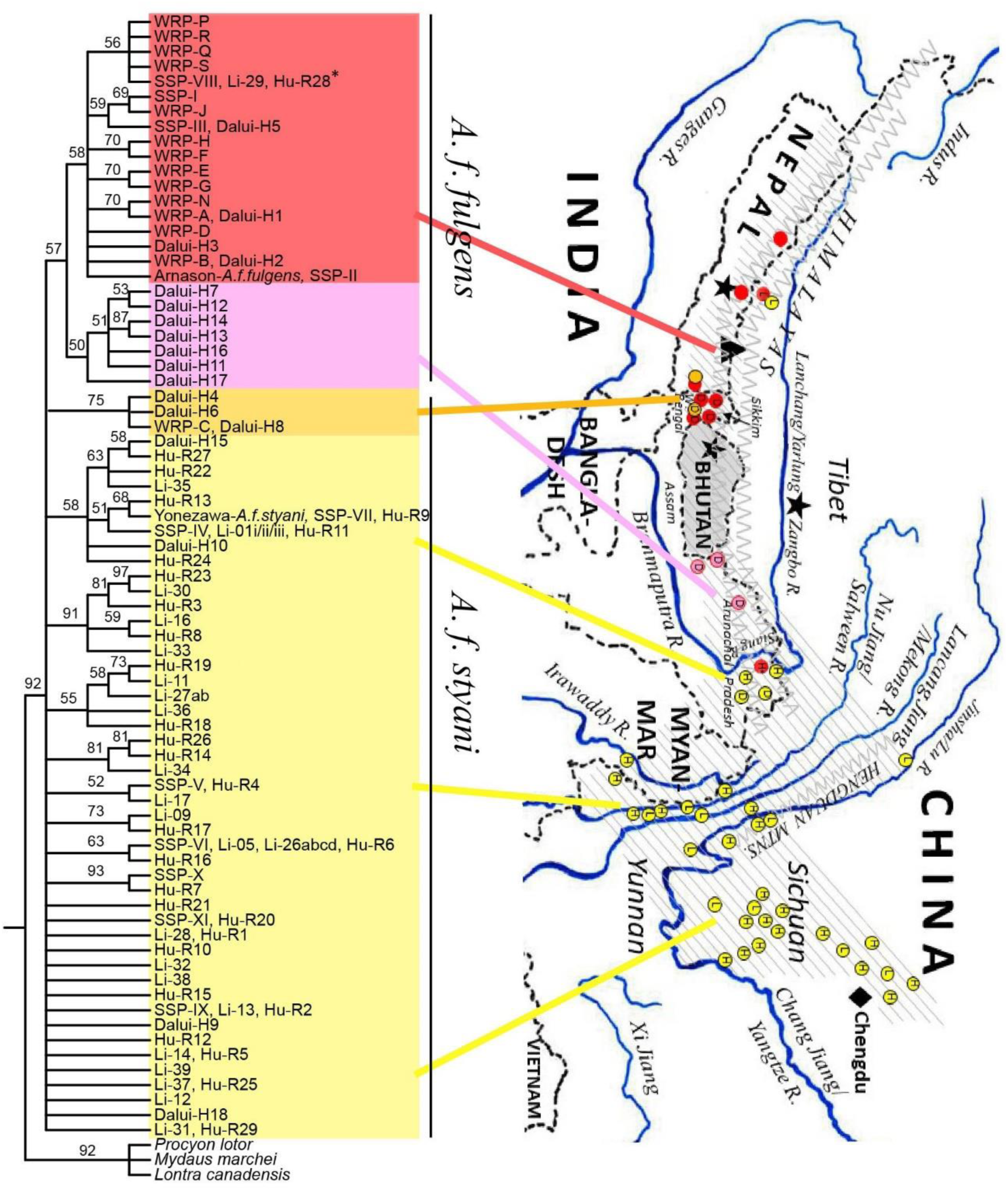
**-left** Maximum Likelihood (ML) cladogram (best tree collapsed) of evolutionary relationships among all known red panda haplotypes (*h* = 75) from five studies^a^ plus two NCBI reference samples of both ‘subspecies’; single representatives from three nearest families to Ailuridae included as outgroups; topology based on 346 positions of mtDNA-CR shared by all 78 haplotypes; bootstrap support (measure of replicate trees in which associated taxa cluster together) denoted in percentages on the branches; phylogenetic analysis conducted in RAxML via the CIPRES portal; **-right** map of south-central Asia depicting presumed range of red pandas at time of studies, with sampling locations for the five studies^a^ denoted on it, and lines drawn to phylogeographic “haplogroups” resulting from sampling in these locations ^a^ L = Li et al. (2005), H = Hu et al. (2011), D = Dalui et al. (2021), [blank] = Dueck & Steffens (2022), Dueck (2021)

One of the small *A. f. styani* subclades in the ML tree contained the three Dalui et al. (2021) subclade 1c haplotypes which they had classified as *A. f. fulgens*, but here confirmed to be affiliated instead with *A. f. styani*, with one haplotype again matching WRP-C. Members of this moderately supported subclade (75%, “*Afs*-orange”), were sourced to West Bengal, eastern Sikkim, and eastern Nepal, west of Bhutan in a region where *Aff*-red was likewise found (**Fig. 2-right**).

The remainder of the *A. f. styani* haplotypes (“*Afs*-yellow”) in the ML tree were all sourced from east of the Siang River (eastern Arunachal Pradesh, northern Myanmar, Yunnan/Sichuan in China, SSP - US zoos, Yonezawa reference), except one (Li-31) with conflicting source information^2^. They were either grouped into eight *A. f. styani* subclades consisting of 2-9 haplotypes with 52-93% support, or were 16 unpaired haplotypes. There was no consistent phylogeographic structure associated with *Afs*-yellow.

Notably, **Fig. 2** demonstrates a “criss-cross” of purported subspecies on either side of Bhutan, with eastern *Afs*-orange on the west side (sympatric with *Aff*-red) and western *Aff*-pink on the east side, but no samples from Bhutan. Additionally, the purported subspecies were minimally sympatric elsewhere – eastern Tibet/western Arunachal Pradesh (*Aff*-red Hu-R28 in *Afs*-yellow territory), and perhaps in south-central Tibet north of Kathmandu, if the source location for Li-29 and Li-31 is correct to Zhangmu.

Another ML phylogenetic tree (not shown), using only the midrange exclusion haplotypes (*h* = 60) and three outgroups provided similar results to the midrange inclusion ML tree of **Fig. 2**, except that *Aff*-red was more strongly supported (86%), while clustering of *Afs*-yellow together remained unsupported.

Phylogenetic analysis of the final dataset with three outgroups using the BI method (not shown) fully supported a monophyletic clade of all red panda haplotypes (1.00 PP) with an almost identical topology to that of the ML tree for the same dataset, except that *Aff*-pink (from east of Bhutan) did not have separate support to cluster within a weakly supported (0.55 PP) *A. f. fulgens* subclade. However, *Aff*-red did have stronger support (0.69 PP) within *A. f. fulgens*, and *Afs*-orange (from Nepal-Sikkim-West Bengal) likewise had stronger support (0.98 PP) within scattered and unclustered *A. f. styani* in the BI tree.

Network analysis of the condensed red panda taxa (**Fig. 3**) also indicates the same grouping as the ML tree of *A. f. fulgens* into two haplogroups (red and pink ovals) with a homoplasy within each and the most contraction for taxa found in Nepal, West Bengal, and Sikkim (red oval), suggesting limited differentiation. Only three mutational sites separate the closest haplotypes of each purported subspecies – Dal-17 (from west Arunachal Pradesh) and Li-31 (from central Tibet). The haplogroup of *A. f. styani* from seven animals sampled in eastern Nepal, eastern Sikkim, and West Bengal (orange oval) is on a long branch extending from other *A. f. styani* found in north Sichuan. Several homoplasies are evident in the rest of *A. f. styani* (yellow oval) from China and Myanmar with up to five mutational steps between some haplotypes and smaller contractions of taxa into similar haplotypes.

**Fig. 3.**
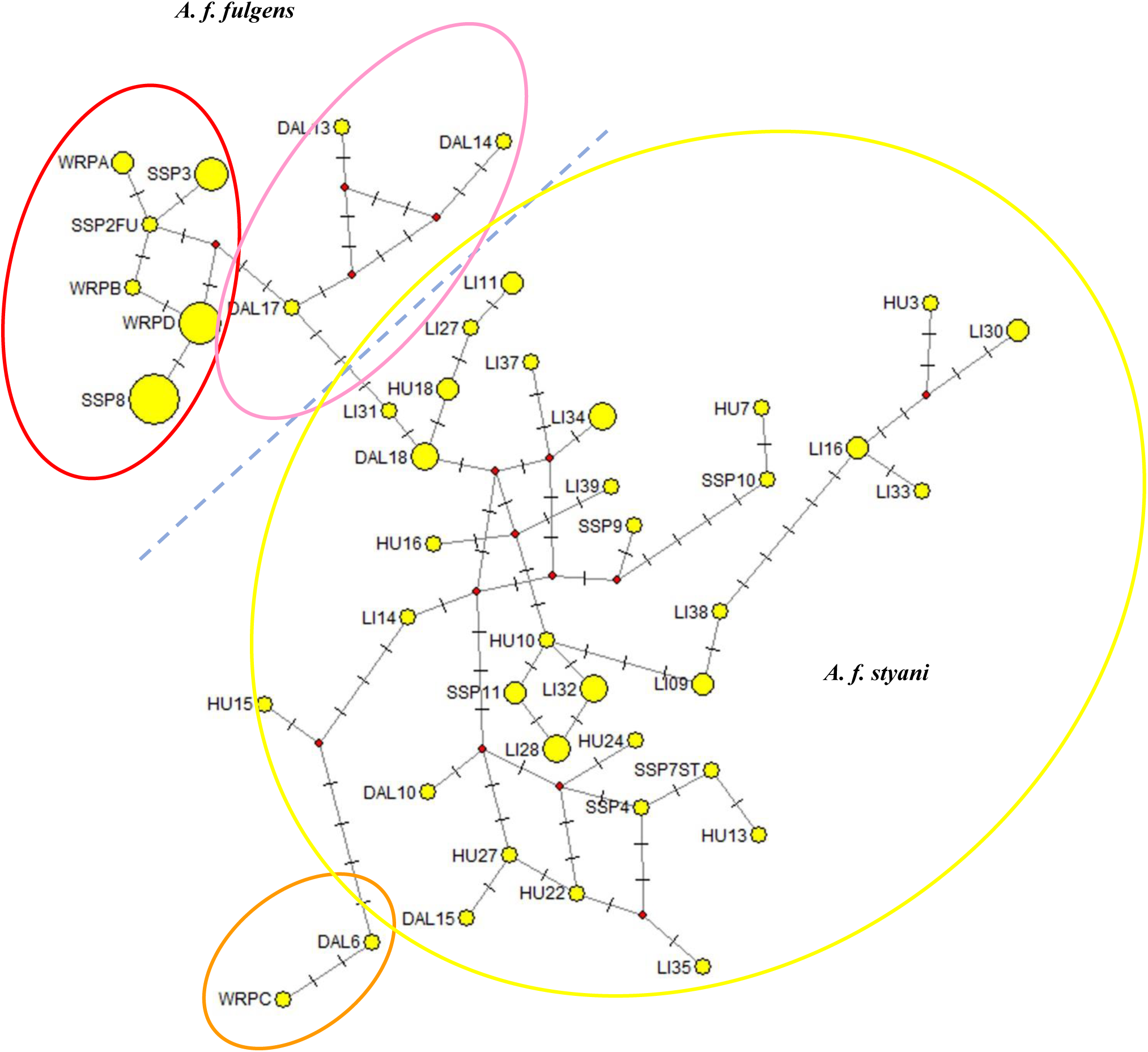
Median-joining network of 75 red panda taxa condensed to 44 haplotypes in star contraction option, calculated in Network 10.2.0.0; haplotype frequencies (number of closely related taxa represented by haplotype) in proportion to size of yellow nodes; small red dots represent median vectors, mutated positions shown by small lines perpendicular to network connectors; homoplasies from reversals or parallel mutations shown by multiple connections between haplotypes; dashed blue line added to indicate separation between purported subspecies, and oval colors group taxa into same haplogroups as in ML cladogram (**Fig. 2**)

Graphs comparing the haplotype diversity and frequency distribution between the two subspecies (**Fig. 4**) illustrate that each has two dominant haplotypes (≥ 12%), but the remainder of *A. f. styani*’s diversity (*n* = 170) was subdivided into eight medium-frequency haplotypes (3-6%) and many (37) low-frequency (< 2%) haplotypes. Alternately, the remainder of the smaller *A. f. fulgens* dataset (*n* = 101) was comprised of six somewhat-larger-category medium-frequency haplotypes (6-9%) and fewer (16) low-frequency (≤ 2%) haplotypes. There were no shared haplotypes between subspecies.

**Fig. 4.**
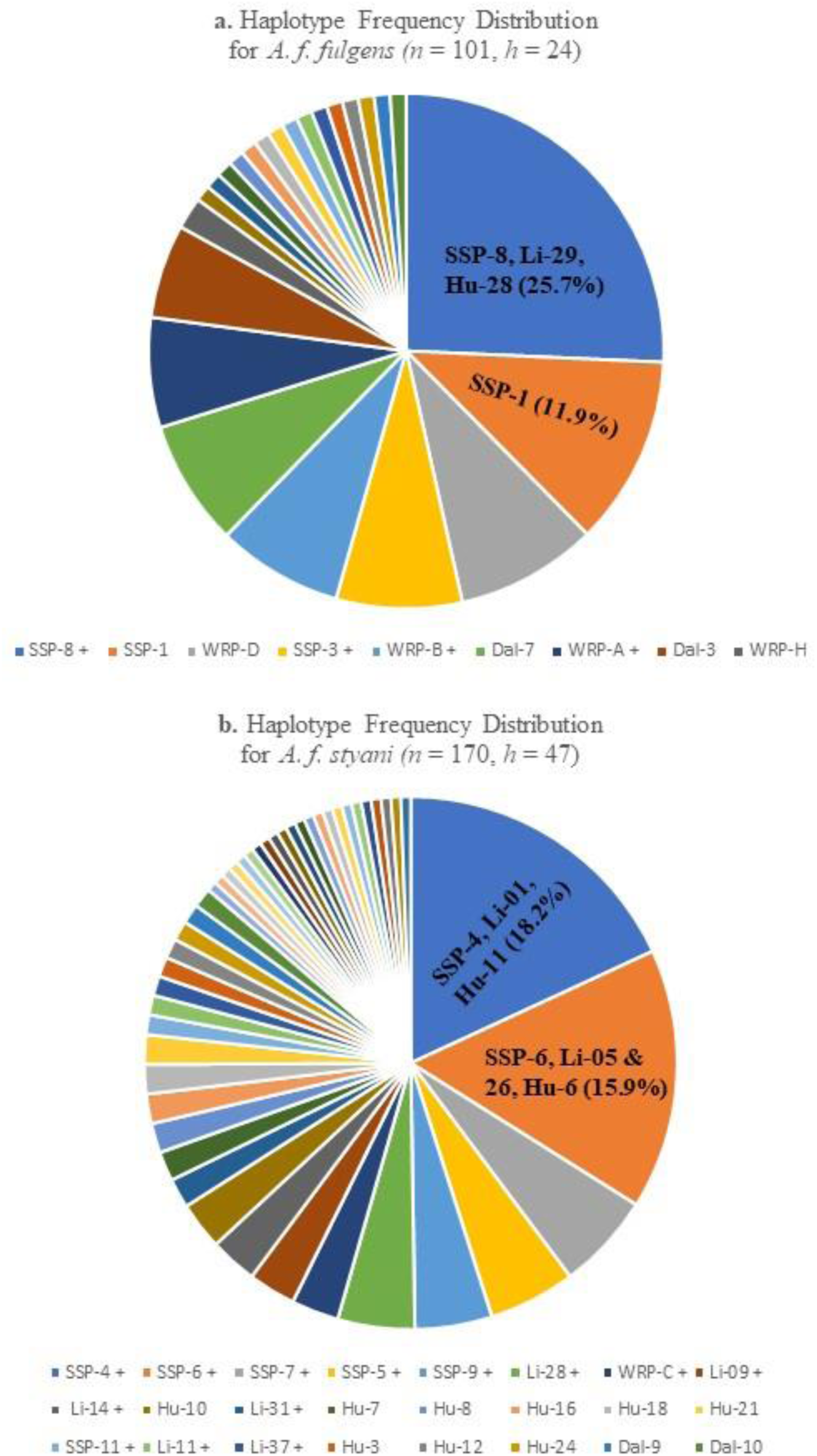
Frequency of mtDNA haplotypes exclusive to each of two ‘subspecies’ of red pandas from native ranges: **a.** *A. f. fulgens* and **b.** *A. f. styani*; assembled from four datasets of previous studies (*N* = 271, *h* = 71); haplotypes in frequencies ≤1.0% not labeled

Statistical estimates of evolutionary divergence based upon pairwise analysis among the 71 haplotypes for native red pandas as a group and divided into two subspecies showed mean pairwise sequence divergence values of 2.65% overall, 3.18% between subspecies, 1.48% for *A. f. fulgens*, and 2.39% for *A. f. styani*.

Population genetic diversity indices for the two subspecies and the four haplogroups from the ML tree are presented in **Table 1**. <u>Haplotype diversity</u> (*Hd*) was high for all red pandas combined (0.96), but marginally higher for *A. f. styani* (0.93, *n* = 170) than for *A. f. fulgens* (0.89, *n* = 101). Likewise, *Afs*-yellow (*n* = 163) had a higher *Hd* (0.92) than *Aff-*red (*n* = 87, 0.86). However, the smaller haplogroups of each subspecies differed in that *Hd* for *Aff-* pink (*n* = 14) was slightly higher (0.60) than *Afs-* orange (*n* = 7, 0.52). <u>Nucleotide diversity</u> (*π*) was 0.0237 for all red pandas combined, with *A. f. styani* having about double that of *A. f. fulgens* – 0.0207 and 0.0096, respectively. Between the larger haplogroups of each subspecies, *π* of *Afs*-yellow was higher (0.0201) than *Aff*-red (0.0078), while *π* in the smaller haplogroups of each subspecies was very similar (*Aff*-pink – 0.0049, *Afs*-orange – 0.0045). <u>Other</u> measures of diversity – *S* (the number of segregating/polymorphic sites), *k* (the average number of nucleotide differences between sequences), and Watterson’s estimator *ϴ_w_* per site and per sequence (population mutation rate) – demonstrated similarly higher values for *A. f. styani* than for *A. f. fulgens*.

**Table 1.** Genetic (mtDNA) diversity indices as calculated in DnaSP6 (DNA Polymorphism), for native red pandas from four studies combined – Li et al. (2005), Hu et al. (2011), Dalui et al. (2021), and Dueck and Steffens (2022).

| Grouping: | ALL | *A. f. fulgens* | *Aff*-red | *Aff*-pink | *A. f. styani* | *Afs*-orange | *Afs*-yellow |
| --- | --- | --- | --- | --- | --- | --- | --- |
| *n* (sequences): | 271 | **101** | 87 | 14 | **170** | 7 | 163 |
| *n* (sites): | 338-9* (340) | **339\* (340)** | 339* (340) | 339* (340) | **338-9\* (340)** | 339* (340) | 339* (340) |
| Variable: |  |  |  |  |  |  |  |
| # haplotypes (*h*) | 61 (71) | **23 (24)** | 17 (17) | 7 (7) | **41 (47)** | 3 (3) | 38 (44) |
| Haplotype diversity (*Hd*)<br>+/- S.D. | **0.955**<br>0.005 | **0.890**<br>**0.018** | 0.861<br>0.023 | 0.604<br>0.150 | **0.928**<br>**0.011** | 0.524<br>0.209 | 0.923<br>0.012 |
| Nucleotide diversity ($\pi$)<br>+/- S.D. | **0.0237\***<br>0.00037 | **0.0096\***<br>0.00060 | 0.0078*<br>0.00043 | 0.0049*<br>0.00186 | **0.0207\***<br>0.00056 | 0.0045*<br>0.00194 | 0.0201*<br>0.00055 |
| Nuc. div. ($\pi$ Jukes-Cantor) | 0.0244 | **0.0096** | 0.0079 | 0.0050 | **0.0201** | 0.0045 | 0.0195 |
| Mean # nuc. diffs.<br>between sequences (*k*) | 8.015* | **3.240\*** | 2.647* | 1.670* | **6.994\*** | 1.524* | 6.781* |
| # segregating sites (*S*) | 56* | **28\*** | 18* | 9* | **39\*** | 4* | 36* |
| Theta ($\Theta_w$) / site | **0.0267\*** | **0.0159\*** | 0.0105* | 0.0084* | **0.0202\*** | 0.0048* | 0.0187* |
| Theta ($\Theta_w$) / sequence | 9.066* | **5.398\*** | 3.573* | 2.830* | **6.831\*** | 1.633* | 6.353* |

**Note:** Values in ( ) are input values. When program **excluded** sites with gaps/missing data, values shown were used/obtained, but that collapsed some haplotypes. If pairwise-deletion and column-by-column options were available, values with \* were used/calculated, which excluded gaps/missing data **only** in pairwise comparisons.

**Table D, Online Resource I,** presents pairwise population differences between subspecies and haplogroups as *F_ST_* and *χ^2^ P*-values. All *χ^2^ P*-values were significantly different (*P* < 0.0001) except *A. f. fulgens* vs *Aff*-red and *A. f. styani* vs *Aff*-yellow (both *ns*), but those haplogroups made up the majority of the subspecies in which they were members. The two smaller haplogroups of each subspecies (*Aff*-pink *vs Afs*-orange) were slightly less significantly different (0.001<*P*<0.01). The *F_ST_* between subspecies was 0.52.

Population demography (size change and stability) was assessed and demonstrated some differences between putative subspecies. Results for tests of neutrality of mutations for subspecies, haplogroups, and combined are given in **Table E, Online Resource I**. <u>Tajima’s *D*</u>, which tests whether all mutations are selectively neutral, was calculated using both *S* and *Eta*. All groups had small negative values (more so for *A. f. fulgens* but none below -2), although results were only significant (*P* < 0.05) for *Aff*-pink using *Eta*. <u>Fu’s *F*</u>*<u>_S_</u>*, which is based on the haplotype frequency distribution conditional on the value of *ϴ_w_*, had larger negative values (more so for *A. f. styani* and especially combined) and thus all were significant except for the two smaller haplogroups (*Aff*-pink and *Afs*-orange) within subspecies. <u>Mismatch distribution plots</u> (not shown) for both subspecies and combined all exhibited the same pattern of changing (*vs* constant) population size. Another simple measure to infer demographic history is by <u>comparing nucleotide diversity measures</u> *π* and *ϴ_w_* per site (highlighted yellow and green, respectively, in **Table 1**). This comparison showed a 13% higher value of *ϴ_w_* for combined and a 66% higher value of *ϴ_w_* for *A. f. fulgens*, representing population size change, but only a 2% higher value of *π* (i.e., almost equal) instead for *A. f. styani*, representing population stability.

The distribution of genetic variation as assessed in AMOVA for both subspecies and haplogroup comparisons (**Tables Fa** and **Fb**, respectively, **Online Resource I**) indicated nearly equal values for between/among and within comparisons. Specifically, 48% of the total variance was found between subspecies and 52% within subspecies. Similarly, 52% of the total variance was found among haplogroups and 48% within haplogroups. The fixation index of differentiation (*F_ST_*) was highly significant (*P* = 0.0000) in both the former (0.48) and latter (0.52) comparisons,^3^ but no haplotypes were shared between subspecies or haplogroups.

## Discussion

We evaluated a comprehensive dataset from all previous red panda mtDNA-CR studies (*N* = 393, *h* = 126) to assess phylogeographic structure from more of their native range than any single study before, filling in some missing links, but also elucidating the complex nature of genetic variation along the geographic arc of red panda distribution.

Our phylogenetic analyses, based on more evenly distributed sampling, several reference standards, and outgroups from two or three families, demonstrates that red pandas comprise only one species, *Ailurus fulgens*, with some phylogeographic patterning (haplogroups) and possible distinction of one sublineage (western *A. f. fulgens*) within, but no sharp boundary between two distinct taxa. Exploratory NJ analyses with and without midrange haplotypes clearly illustrated the impact that narrowing the sampling gap had on improving taxonomic accuracy of species delimitation for red pandas. Our network analysis showed there are more mutational sites between some eastern (*A. f. styani*) haplotypes than between haplotypes of different subspecies. In the ML cladogram, there is minimal support for the clustering of *A. f. fulgens* haplotypes, thus it acted as a larger, more complex subclade within the species rather than a separate entity, since the *A. f. styani* did not likewise cluster together. Bayesian methodology confirmed the ML tree topology with only a few minor differences in support. It is therefore apparent from our phylogenetic analyses of a more broadly sampled mtDNA-CR dataset that the arbitrary division between red pandas is nothing more than geographic variation within a single, strongly supported monophyletic clade – i.e., red pandas are one species having a single common ancestor.

Equally important, the map associated with the ML cladogram plotting sample origin demonstrates sympatry and a possible cline in the midrange of red panda distribution between eastern and western haplogroups, suggesting ongoing gene flow across a contact zone between so-called ‘subspecies’. For instance, one haplotype found in two individuals from eastern Nepal in 2003 (Dueck and Steffens 2022) was also discovered *circa* 2020 in three individuals from India bordering Nepal, as well as single individuals of two other haplotypes that clustered with it (Dalui et al. 2021); all three haplotypes associated with *A. f. styani* (*Afs*-orange) but were found together west of Bhutan, in previously assumed *A. f. fulgens*-only territory. Several hypotheses are offered for this distribution: 1) transplantation, except zoo reintroductions occurred after the first incidence of “errant” *A. f. styani* (Jha 2011); 2) sustained natural incursion across the contact zone; 3) relictual and isolated self-sustaining enclave of *A. f. styani* with some innate reproductive barrier; and 4) persistence of *A. f. styani* haplotypes via matriarchal inheritance from previous incursion in a mixed population with no reproductive barriers, continuous for at least three generations. A few other occurrences of sympatry were discovered as well, and some clinal variation seemed apparent from presence of *Aff*-pink, a subset of western *A. f. fulgens* but phylogenetically intermediate to eastern *A. f. styani*, just east of Bhutan.

There has been considerable debate as to what physiographic boundary (e.g., Yangtze, Nujiang, or Siang River) separated the ‘subspecies’, but our genetic evidence suggests there may be no boundary, nor strict division, between western and eastern sublineages of red pandas in their native range. However, genetic evidence to address these hypotheses from the potential contact/hybridization zone in Bhutan is still sorely lacking, and nominal from northern Myanmar, but for different reasons. Genetic studies have not been permitted in Bhutan, and access to research areas in Myanmar is both difficult and dangerous (further discussed in **Online Resource II**).

Population genetic analyses indicated that both differentiation indices (*F_ST_* and *χ^2^ P*-values) between ‘subspecies’ and between haplogroups were significant, as expected when no haplotypes were shared despite being very similar. Furthermore, AMOVA results show as much variation within as between/among groups. Evolutionary divergence estimates based on pairwise comparisons showed that haplotype sequence divergence was just 0.53% greater between ‘subspecies’ than among all 71 haplotypes assessed, another indication of little separation between ‘subspecies’ and close relationships among haplotypes. Nucleotide diversity (*π*) values were high overall (0.024), but the western *Aff* sublineage had only about half that of the eastern *Afs* group, meaning that the *Aff* sublineage was more closely related, noticeable from the network diagram and lower divergence value. By analyzing correlates of *π* for a dataset of 639 mammalian species, James and Eyre-Walker (2020) determined that *π* was significantly correlated to range size, and species with larger range sizes have larger effective population sizes. **Fig. 2**’s map demonstrates that the western sublineage may have a smaller range size than the eastern group, thus a lower *π* might be expected. Interestingly, haplotype diversity (*Hd*), also high overall (0.96), was nearly the same between ‘subspecies’. This is surprising because there were two-thirds more samples and twice the number of haplotypes in the eastern *Afs* group than the western *Aff* sublineage, so a new haplotype was encountered more frequently in the eastern (1/3.6) than western (1/4.2) groups, and logically more haplotypes are found with more sampling, to a point. But in addition to sample size, haplotype frequency is used to calculate *Hd*, so that must be relatively balanced between the two groups. As depicted in the frequency distributions of **Fig. 4**, both ‘subspecies’ groups had relatively even, genetically healthy allocations of common, moderate, and rare haplotypes given differing sample sizes. In a broad survey of mtDNA diversity among mammals, Saitoh (2021) found a mean *Hd* of 0.578 for 66 terrestrial species, so native *Ailurus fulgens* exceeded that. A reminder, however, is that our dataset was compiled from samples collected over 20 years and may not reflect current genetic diversity status.

### Demography, biogeography, and geographic variation

The four simple measures of demographic history we presented suggest some population size change of red pandas overall, as generally shown by the mismatch distributions. Both tests of neutrality of mutations exhibited negative values from an excess of low frequency mutations, indicating population expansion as after a bottleneck or selective sweep. Results from *Fu’s F_S,_* Tajimas’s *D*, and comparison of nucleotide diversity measures produced contradictory results for which ‘subspecies’ had more change or more stability. Otherwise, it is difficult to draw specific conclusions from our demography analysis other than relatively recent polymorphisms indicate some population expansion in all native red pandas. We do not wish to speculate further about demographic history with our combined dataset and limited analyses as we believe there is too much leeway given debatable calibration bases and broad confidence limits in such a short time frame which might lead to questionable conclusions. Fu and Wen (2023) were also skeptical about over-reliance on timing of phylogenetic-geologic correlations based on sequence data due to time estimation being dependent on so many variable factors.

However, fossil records indicate that advanced red panda predecessors appeared during the mid-Miocene as a smaller hypocarnivore lineage with a Holarctic distribution (Wallace 2011), but extant red pandas likely diverged from ailurine ancestors in Asia about the time of Pleistocene onset ∼2.5mya (Li et al. 2005). This epoch is the earth’s most recent period of long glaciations and shorter interstadials over much of the Northern Hemisphere. It caused significant north-south displacement of temperate-zone biota, often leading to multiple re-distributions of genetic variation in animal populations that had contracted and mixed in refugia (Hewitt 2000). However, things were different in continental East Asia with its unique geography, which had less ice coverage but asynchronous glaciations on the Tibetan Plateau. There were still strong climatic oscillations overall to which vegetation responded with range contraction and expansions, but its Pleistocene climate was comparatively mild. Glacial refugia were numerous, expansion differed greatly in time, scale, and direction, and impacts on species history varied drastically but were felt less in the southwest. Phylogeographic patterns, highly consistent between plants and animals, were therefore more complex. Organisms often retained older genetic histories associated less with climatic than with geologic events such as plateau upheaval, development of the three-step terrain in southwest China, and fluvial changes (Fu and Wen 2023). Interestingly, in the high mountains of the Himalaya, fluctuations in glacial coverage during the Pleistocene indicate that the early local Last Glacial Maximum (LGM) of 60-27kya was the most severe, while the global LGM (18-24kya) was very restricted in extent, generally not exceeding 10 km from current ice margins due to lower insolation, weaker monsoon, and less snow accumulation at high altitudes; modest glacial advances were merely from reduced temperature (Owen et al. 2002).

How may have these environmental changes impacted the genetic diversity and structure of native red panda populations? First, geographic variation in many attributes is anticipated of organisms inhabiting a long narrow range, especially one bordering different mountain systems, fragmented by rivers, and historically impacted by climatic fluctuations (Endler 1977; Hillis 2020). For the relatively recently evolved red panda during the severe early glaciations of the Pleistocene, perhaps a more homogeneous, widespread, panmictic population developed as it converged on numerous scattered refugia and expanded multiple times, following Hewitt’s (2000) pioneer model of genetic re-distribution. But the last glacial episode had a relatively insignificant impact on mobile biota in Himalaya (Fu and Wen 2023), likely more vertical than lateral^4^, and red pandas are considered “altitudinal migrants” anyway (Glatston et al. 2015), following bamboo availability and climatic optimums. Less disruption may have allowed the early geographic variation present in western populations to develop sublineage structure, semi-isolated yet interconnected via metapopulations as Dueck and Stephens (2022) demonstrated for regional structure in Nepal, while the eastern populations maintained more homogeneity and stability via increased gene flow.

Nonetheless, the sympatric presence of both ‘subspecies’ in at least one region through time demands further investigation. Although the ‘errant’ *Afs*-orange haplogroup appears intermediary between ‘subspecies’ in the NJ phylogram, thereby shrinking the gap and implying a cline, network and BI analyses indicate a more distant relationship, related more to samples taken at the northeast extreme of the range than the middle, suggesting it reflects an older signature of distribution from a common source and/or early glacial re-shuffling maintained *in situ*. However, since no empirical evidence of any intrinsic barriers between ‘subspecies’ is known, there is also no reason to believe that they are not now merging, given this apparent gene flow in the midrange. To determine whether the lineages are evolving independently, if this is an example of parapatric divergence, or if intraspecific variation follows a cline, their contact zone must be intensively sampled to search for evidence of reproductive isolation (assortative mating, lower production or fitness of “hybrids”). If none is found, then likely the widely distributed subpopulations of red pandas are part of a geographically variable, occasionally locally interbreeding, and single evolutionary lineage (Chambers et al. 2023).

### Limitations of mtDNA

Some caveats should be noted about total reliance on the mtDNA genome, effectively a single locus. While mtDNA was especially valuable for phylogenetic reconstruction leading to the field of phylogeography, its maternal inheritance and stochastic variability between loci limits its utility (Allendorf 2016). Haplogroups cannot be expected to reflect historical biparental lineage (Hillis 2020), although White et al. (2008) documented some exceptions to this tenet. We only used a small segment of the hypervariable CR upon which to base our conclusions, likely underestimating the genetic diversity present (Banes and Galdikas 2016). Multiple recurrent mutations at the same site are not uncommon, and that instability may misinform about true differences (Avise 2004). MtDNA may be subject to incomplete lineage sorting (ILS) when retention of ancestral polymorphisms confounds congruity with species trees from nuclear genes, but may better reflect a species tree in shorter-term if lineage sorting is complete (Moore 1995). However, ILS was recently found to be only a minor explanation for phylogenomic conflicts among genes (Scornavacca and Gaultier 2017). Inconsistencies with patterns of nuclear gene flow may occur from reproductive interactions between lineages via hybridization (Hillis 2020). Differential introgression can obscure species history by introducing potential bias for inferring demographic properties or evolutionary histories, leading to discordance with gene trees from nuclear DNA (Harrison 1989; Ballard and Whitlock 2004; Marshall et al. 2021). Mito-nuclear discordance could also be caused by small effective population size or sex-biased dispersal (Ballard and Whitlock 2004). To further complicate matters, mtDNA was traditionally considered to be selectively neutral and unlinked to the nuclear genome, but some studies suggest that both direct and indirect selection influence mitochondria, given its pivotal role in bioenergetics (Dowling et al. 2008; Gershoni et al. 2009; Ballard and Melvin 2010). And while there is often a problem in defining species boundaries with mtDNA due to oversplitting (Hillis 2020), this was not the case in our study.

Yet mtDNA remains a valid, valuable indicator of phylogeographic patterns for rare, elusive, endangered species. It is the first step to resolve recently isolated groups, having a more appropriate timescale for coalescence than nuclear loci (Barrowclough and Zink 2009; Marko and Hart 2011). For instance, Hung et al. (2015) compared results from mtDNA and nuclear loci in birds and demonstrated that mtDNA was indeed robust at estimating inter-population divergence, especially for species with recent evolutionary history. In fact, mtDNA sequencing was the only genetic tool used for phylogeographic inference together with ecological niche modeling and fossil data to track giant panda history and distribution during the LGM (Luna-Aranguré and Vázquez-Domínguez 2021). However, to most accurately delimit species and assess gene flow over space and time, mtDNA should be combined with nuclear data (Marko and Hart 2011), as well as morphological, ecological, behavioral, and cytogenetic information as part of an integrated approach (Rubinoff and Holland 2005). Nonetheless, for our study assessing red pandas as thoroughly across their range as possible, it was the only comparable data available.

### Concerns with phylogenomic study declaring red pandas as two separate species

Hu et al. (2020) undoubtedly advanced the molecular understanding of red pandas with their genomic study, especially in identifying nuclear loci that affect fitness. But their taxonomic proposition to separate *Ailurus* into two species, which could have significant conservation implications for an endangered animal, was based on seriously incomplete sampling of red pandas throughout their native distribution. In particular, they left out the entire midrange between the two extremes of the distributional arc, fully one-third of prime red panda territory (850-950km of the ∼2500km-long range, from the eastern Nepal-India border, through Bhutan and Arunachal Pradesh, across the Siang River to southeastern Tibet). Instead, after only sampling in Nepal, Yunnan and Sichuan in China, and minimally in southeast Tibet and Myanmar, they inferred the Yalu Zhangbu/Siang River as the definitive border between two phylogenetic species of red panda, without including any samples from northern India or Bhutan. The omitted territory was precisely the most critical geographic area to sample intensively for taxonomic decisions because of its optimal habitat in the contact and/or potential hybridization zone between the putative subspecies. More informed results from sampling contact zones and into potentially parapatric ranges can distinguish intraspecific geographic variation of populations isolated by distance from interspecific lineages isolated by reproductive barriers, and test the evolutionary independence of populations that exchange genes (Hillis et al. 2021; Marshall et al. 2021; Chambers et al. 2023). Experimental design can have major impacts on the outcome of a study when more emphasis is given to the genetic analyses employed than the sampling strategy; omitting a crucial population segment can bias the other results, whereby gaps in sampling can lead to a false signal of clustering when there is actually only an isolation-by-distance pattern present (Meirmans 2015). Any taxonomic change re-defining a species, especially endangered ones, should be offered as a conservative hypothesis based upon the best and most complete information available, and not be influenced by partisan agendas (Hillis 2022). Anything less does a disservice to the species by putting its management in jeopardy, and to society by creating a misunderstanding of its basic biology.

Several other issues were troublesome about Hu et al.’s (2020) study. First, they only used one outgroup (ferret) to root all three phylogenetic analyses. A single outgroup does not provide support for the ingroup and is likely to misidentify the root position; additional outgroups, either more from the same sister taxa or different taxa, increases the chance of obtaining correct tree topologies, which informs on the directionality of evolution (Smith 1994). In the case of the taxonomically isolated red panda, multiple Musteloidea should be included since outgroup choice can influence support strength of ingroup topologies from possible long-branch attraction problems, generating non-phylogenetic signal (Bergsten 2005; Flynn et al. 2005; Phillipe et al. 2011).

Second, Hu et al.’s (2020) declaration of separate species was reliant upon choice of the Phylogenetic Species Concept (PSC), which does not recognize subspecies. Under the PSC, “species are the smallest, irreducible, monophyletic units as measured by molecular markers” (Haig et al. 2006), combining qualifications of monophyly and diagnosability (Zachos 2016). However, its application, recently popular in mammalian taxonomy, is prone to over-splitting, causing taxonomic inflation (Zachos 2018). Logically, if only the range ends of a geographically variable species are sampled, the PSC might well split up that species. But the red panda distribution is a continuum between range ends with semi-isolated populations throughout, differentially subjected to historical migration as well as mutation, selection, and drift. There are recognizable indications of some phylogeographic structure and gene flow over this range that should be acknowledged and maintained within red pandas to preserve specific adaptations and evolutionary potential. Frankham et al. (2012) proposed that choice of concepts to define a species can have major impacts on the ability to conserve it, and that the PSC was “unsuitable for use in conservation contexts”. Minimum costs and maximum benefits to fitness accrue when reproductive isolation is considered in a species definition such as the Biological Species Concept (BSC). Since species boundaries are by nature “fuzzy" and semi-permeable (Harrison and Larson 2014), distinction may better be diagnosed by minimal gene flow and reduced reproductive fitness in hybrids (Frankham et al. 2012). This idea of incorporating reproductive barriers to gene flow has resurfaced in delimiting phylogeographic lineages as species or subspecies even after genomic or mtDNA evidence suggests divergence as a way to describe more biodiversity without adding more species (Dufresnes et al. 2023). Vences et al. (2024) clearly outline methods to quantify lineage compatibility in hybrid zones for this purpose. Fortunately, the rapid paradigm shift from the BSC to the restrictive PSC from advances in molecular technology is relaxing, with broader consideration towards defining conservation units, such as the above hierarchical system of species>subspecies>populations, a species-population continuum (Coates et al. 2018), and phylogenetic tree-thinking *vs* taxon ranking (Matzke 2023).

The third concern with Hu et al.’s (2020) study involves specimen under-utilization. It is unfortunate that they neglected to collect, or publish if collected, any morphological information about each of the 81 animals captured in the wild whose DNA was used differentially among three genetic systems. This was the perfect opportunity to correlate several data types by individual instead of relying on Groves’ (2011) small sample size of museum specimens and others’ photos to define morphological data supporting their conclusion of separate species^5^. Bista et al. (2021) captured wild red pandas and collected multiple data types with improved techniques that considered animal welfare and field assistant safety. It is also curious that Hu et al. (2020) did not use the same number of samples of mtDNA and nuDNA for an equitable comparison.

Conversely, they did control for sample size (*n* = 49) and (sub)species distribution in comparing mtDNA and Y-chromosome results. But the fourth concern is that they concluded male philopatry and female-biased dispersal in red pandas based solely on population structure results from these uniparentally inherited markers. In contrast, Dueck and Steffens (2022) were able to discern some matriarchal structure here and in a previous Nepal study. However, ascertaining the direction of sex-biased dispersal is more complex. On the genetic side, some studies have found discrepancies between lineage distributions for mtDNA and the Y-chromosome, topology incongruence, and unequal rates of evolution (Boissinot and Boursot 1997; Tosi et al. 2000; Nakagome et al. 2008). On the behavioral side, social systems were found to play a key role in dispersal. Since dispersal applies to the behavior of sub-adults moving away from their natal area, integration of molecular and observational data within a theoretical framework is necessary to understand the evolutionary patterns of gene flow that have developed (Lawson Handley and Perrin 2007). An example of a study combining methods demonstrated male-biased dispersal for an average distance of 21 km via radiotracking and female philopatry via genetics in a related musteloid, the Eurasian otter (Quaglietta et al. 2013). Another example in a shark study disproved sex-biased gene flow patterns discerned by mtDNA and microsatellites when electronic tags revealed equal migration but unequal reproductive integration (Allendorf 2016). For red pandas which are solitary except during breeding season, they soon establish their own territory to acquire resources or mates after dispersal. Radiotelemetry studies of red pandas have shown that these home range sizes vary substantially from 0.9 - 5.12 km^2^, with males’ almost double that of and often overlapping several females’ territories, likely due to their polygynous mating system (Bista et al. 2022). Greenwood (1980) hypothesized that dispersal direction was linked to female-defense polygyny, leading to male-biased dispersal, common in mammalian breeding systems. Little is known about parental care in the wild, but zoo cubs in a naturalized setting stay with mothers after birth for a three-month denning period and up to five-month juvenile period following a three-to-five-month gestation period, leaving little time for re-locating given their circannual reproductive cycle (Curry 2022; Gebauer 2022), another reason to expect female site philopatry. Thus, we advise caution in assuming otherwise until further studies using several data types are undertaken.

### Conservation implications

Taxonomy and the dynamic science-based process determining it are critically important to mitigate global biodiversity loss (Thomson et al. 2018). Taxonomy should reflect our best, most complete information about the biology of taxa upon which to base the most useful management strategies for conservation (Hillis 2022). Omitting a significant portion of red pandas’ range from sampling and analyses, particularly the contact zone between two proposed taxa, does not provide adequate information to justify splitting *Ailurus* into two species, no matter the genetic marker used. It would thus be biologically misleading to say that red pandas are different species that can be managed independently, especially if there is some evidence of gene flow among them. In fact, the admixture analyses of Hu et al. (2020) demonstrated gene flow among their population groupings. Gene flow is critical for long-term survival of a species connected over such a long geographic range, therefore populations within it cannot be managed as isolated units. Doing so would imply that red pandas are evolving independently on each side of some distinct boundary when they are not. There is likely a reproductive continuum across an extended contact zone, much of which remains unsampled (e.g., Bhutan), that connects them. Any separation into the red panda haplogroups shown in our broad mtDNA survey is completely expected as geographic variation and very minor compared to divergence of ailurids from other musteloids or within other musteloid families, so arbitrary division into two species from the genomic analyses based on the incomplete sampling of Hu et al. (2020) is rather meaningless.

Nonetheless, Hu et al.’s (2020) taxonomic revision of splitting red pandas into two species, popularized by media attention (e.g., BBC, CNN, CBC, The Guardian, Newsweek, PBS stories in 2020) and apparently accepted without challenge, will undoubtedly be a factor in the upcoming 2025 ten-year reassessment of red pandas’ status on the IUCN Red List (Glatston et al. 2022). A decision by the international conservation community will therefore be necessary about whether to accept that revision, or to retain the current taxonomy of one species advocated here. Red pandas are currently accepted by the Red List as one species, *Ailurus fulgens*, although a taxonomic note added that Groves (2011) suggested the two taxa (i.e., subspecies) of Ailuridae be treated as separate species.

Technically, a decision to embrace the two-species revision should have no impact on the Endangered Status under which all red pandas are listed, since taxonomic revision (e.g., species splitting) is not considered a genuine reason for changing threat category in the Red List (https://www.iucnredlist.org/assessment/reasons-changing-category; accessed 3/28/24). It is therefore unclear what, if any, advantage species-splitting would have on red pandas’ IUCN listing, since subspecies assessments (assumed to include independent threat evaluations) can also be accepted as long as the species-level assessment is done. Ranking taxa for conservation priority is presently only dependent on threat category in the Red List (from population size trends, distribution/habitat changes, threats identified, actions required), although more inclusive systems have been proposed elsewhere, including consideration of evolutionary uniqueness (Redding and Mooers 2006; Tims and Alroy 2022), which would elevate the overall status of red pandas substantially.

However, if the proposition of two red panda species is accepted, separate assessments would be required, including changes to underlying databases. It would mean international cooperation to re-organize conservation strategies, management, and legislation, all of which take time that red pandas do not have to avoid extinction. But more importantly, it would cut off necessary gene flow and contact between the newly separated entities *via* independent management strategies. It could prevent or inextricably confound genetic rescue if ever necessary to preserve adaptive potential (Frankham et al. 2012; Hamilton and Miller 2015), such as consolidating captive “ark” populations of red pandas formerly considered subspecies (Powell 2023). It could also mean that American zoos can no longer place single cubs of the two “species” together to learn socialization skills^6^. And ultimately it would be biologically misrepresentative.

Unfortunately, the primary author of the 2015 IUCN assessment and likely involved in the upcoming reassessment, dismisses discussions on differences in species concepts as unhelpful to conservation (Glatston et al. 2022), despite taxonomic accuracy being imperative to decisions on appropriate conservation management. She also speculated on the impact of this species-splitting to conservation, suggesting it did not yet have the boost to conservation expected, since taxonomic instability is seen as a real problem. Species-splitting was actually thought to increase protection, with name changes having least effect on charismatic fauna (Morrison et al. 2009). In fact, recent changes to the ‘boundary’ between (sub)species apparently disappointed stakeholders because with *A. (f.) styani* not solely endemic to China now, their threat category cannot be elevated from threatened to endangered. Red pandas face different threats in different regions of their range, and it is probable that red pandas native to parts of the western range are in more immediate danger of extinction (Glatston et al. 2022). But accommodation of different threat categories is not an appropriate reason to upgrade underlying alpha taxonomy (Hillis 2022); any change can be handled by a subspecies assessment within the main IUCN species listing, although that may only be applicable to *A. f. fulgens*. The small hunted population in Myanmar of the eastern group may also be at more risk (Lin et al. 2022), but could be managed nationally. Fortunately, Nepal and Bhutan have recently initiated conservation action plans for red pandas to enhance protection there. We therefore advocate maintaining the current taxonomy of red pandas as one geographically variable, interconnected species from local gene exchange, with regional risk management and cooperation, so that conservation energy can avert diversion from indecision and regrouping and instead focus on immediate action for preserving as much evolutionary potential as possible.

## Conclusions

Summarizing, our comprehensive mtDNA-CR study demonstrated that red pandas are only one species, but may include a more closely related western sublineage, with further indications of phylogeographic organization, gene flow, and a cline. A recent phylogenomic survey from range ends split red pandas into two distinct species, among other issues. We tested whether omission of midrange samples yielded similar results with our data and some minimally concurred. Therefore, we challenge this taxonomic splitting based upon incomplete sampling as misinforming about their basic biology, thereby jeopardizing appropriate conservation management strategies. We recommend a thorough investigation of the contact zone in the midrange of red panda distribution (Myanmar, northern India, and especially Bhutan) to evaluate intergradation with multiple data types, meanwhile maintaining the current taxonomic rank of one *Ailurus* species. The western sublineage (*A. f. fulgens*) as presently known from Nepal, Tibet, and India surrounding Bhutan could then be acknowledged as a geographic variant by the term ‘subspecies’ for separate assessment within the species framework for an elevated Red List threat categorization. This would allow the IUCN’s 2025 overall reassessment of red pandas to proceed more efficiently towards action, rather than be stymied by bureaucratic difficulties due to taxonomic inflation. A conservative solution would benefit researchers, conservation planners, and especially red pandas, providing more flexibility by recognizing more biodiversity without more species, using the best, most complete information available. It is time we bridge the gap for all red pandas, not burn the bridge between them.

## Supporting information

Suppl Info_Missing links - red pandas

## Acknowledgements

We thank David Hillis, Prof., Dept. of Integrative Biology, U. Texas-Austin, for encouragement, discussion, and advice before writing; Anthony Davis, Prof. Emeritus, Dept. of Geography, U. Toronto, for prior encouragement, comments and discussion during writing; all those associated with the SREL DNA lab and AZA’s Red Panda SSP acknowledged in earlier publications; and previous co-author Erik Steffens.

## Statements and Declarations

### Employment

The first author, Lucy Dueck, was employed at Savannah River Ecology Laboratory when the red panda genetics project and concept of comparison between studies first began, but has since retired.

### Author Contributions

Both authors contributed to the study concept and design. Data compilation, map drawing, NJ/MEGA, Network, and DnaSP analyses were performed by Lucy Dueck; ML, BI, and AMOVA analyses, map-tree compilation, and data cataloging were performed by Deniz Aygoren Uluer. The first draft of the manuscript was written by Lucy Dueck, and Deniz Aygoren Uluer commented on previous versions of the manuscript. Both authors read and approved the final manuscript.

### Data Availability

The haplotype sequences of red pandas used in this study can be obtained in GenBank using accession numbers found in **Table A, Online Resources**. GenBank accession numbers for outgroups are: *Mydaus marchei* AY587074, *Mephitis mephitis* AY587106, *Mephitis mephitis* AY587094, *Mephitis macroura* AY5871051, *Spilogale putorius* AY587075, *Spilogale gracilis* AY587078, *Conepatus leuconotus* AY587093, *Conepatus chinga* AY159818, *Procyon lotor* AB297804, and *Lontra canadensis* = MT513950.

## Dedication

This paper is dedicated to Luna, the oldest red panda (6/22/01-3/25/24) and treasured member of the SSP, whom I met as a cub when I first began this project in the early 2000s and whose survival inspired me to undertake this study, and to my mother, who always asked “But what good is it?” when I explained my work. – L.D.

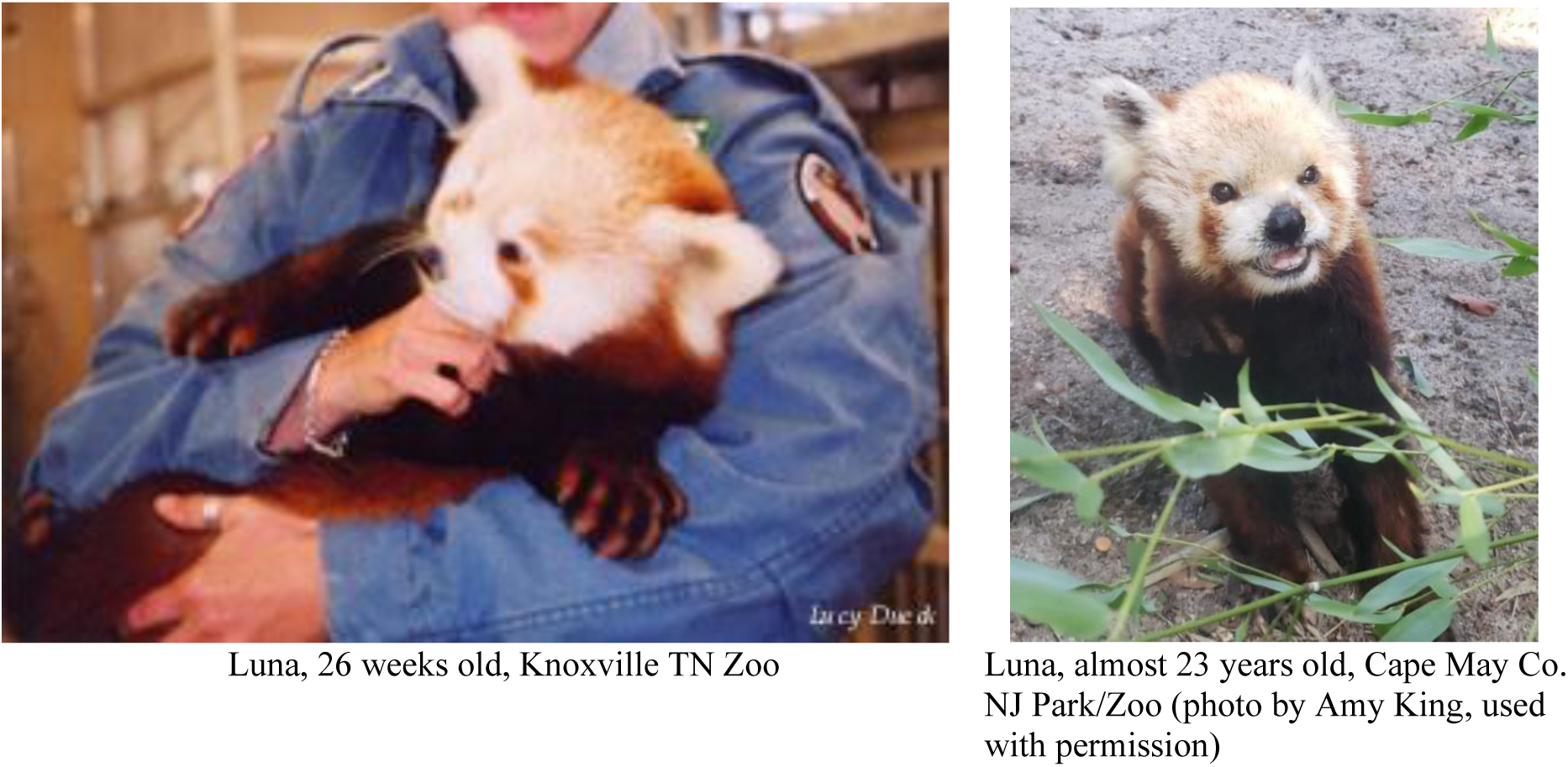

## Footnotes

1 Some site irregularities noted with Li et al. (2005); haplotype location not site-specific in Hu et al. (2011)

2 Zhangmu, E.? Tibet (Li et al. 2005)

3 *F_ST_* values may differ slightly between programs due to usage of positions with gaps/missing data.

4 See Kattel et al. (2013, 2015) for discussion on impact of seasonal wind shift, monsoon, topography, and temperature lapse rate.

5 Interestingly, half of their Table S1 for morphological differences between “species” includes observations made by Pocock (1941) who concluded there was too much variation within each group to justify full species status.

6 See **Photos A** and **B, Online Resource IV,** demonstrating this practice.

