## Supplementary material for "Missing links connect the phylogeographic structure of endangered red pandas, remaining as one species – *Ailurus fulgens*, and expediting conservation": Suppl Info_Missing links - red pandas

#### I. TABLES

**Biodiversity and Conservation**

Lucy A. Dueck[1] and Deniz Aygören Uluer

**Table A.** Summary of comparable mtDNA-CR studies on red pandas (*Ailurus fulgens*) in US zoos and native lands

| **Study author(s)**<br>**Yr. coll./pub.** | **Dueck**<br>**2002/2021** | Su et al. ǂ<br>?/2001 | Dueck & Steffens<br>2003/2022 | Li et al.<br>?/2005 | Hu et al.<br>?/2011 | Dalui et al.<br>2020?/2021 | references<br>?/2007 |
|---|---|---|---|---|---|---|---|
| | * * * * * * * * * * | * * * * * * * * * * | * * * * * * * * * * | * * * * * * * * * * | * * * * * * * * * * | * * * * * * * * * * | * * * * * * * * * * |
| **Sample locations** | USA zoos | China (Sich-uan, Yunnan) | Nepal (Ilam, Lang-tang, Manang) | China (Sich-uan, Yunnan, Tibet), Burma | China (Sich-uan, Yunnan, Tibet), Burma | India (Sikkim, W. Bengal, Aru-nachal Pradesh) | *A.f.f.* - unknown; *A.f.s.* – Jap. zoo |
| **# bp seq'd. in CR** | 383 | 236 | 383 | ~551 | 551 | 436 | all |
| **# bp in common with base dataset** | 383 | 219 | 383 | 383 | 383 | 339 | 383 |
| **Sample size (*N*)** | 67 | 53 | [68 | 40 | 119 | 44] | 2 |
| **# haplotypes** | 11 | 25 | 17 | 24^ | 29 | 18 | 2 |
| **Source (sample)** | zoos (hair/ blood) | zoos (hair/ blood), museums (leather) | wild (fecal) | captured wild (hair), museums (skin) | wild (fecal), zoos/museums (muscle/skin/ blood/hair) | wild (fecal) | *A.f.f.* - unknown; *A.f.s.* - liver/muscle/blood |
| **GenBank accession #s** | MT513939-MT513949 | AF294229-AF294253 | OK381584-OK381597 | AF291579-AF291586, AY849715-AY849736 | HQ992964-HQ992985 (+7 same as some of Li's) | MT891293-MT891310* | Arnason *A.f.f.* = NC_011124; Yonezawa *A.f.s.* = NC_009691 |

NOTES: Burma = Myanmar; ^ = Li 40 (1/25) omitted because too many missing bases; * MT891307 = haplotype 9, not 19;

[-] = samples used for pop. gen. analyses; ǂ = omitted from combined analyses due to short sequence length, but assessed separately.

**Table B.** Mutational sites from consensus sequence for all 75 mtDNA red panda haplotypes used in this study, color-coded to match subclades as shown in ML cladogram (**Fig. 2**). Grey color of position numbers and changes indicate affinity of haplotypes from intermediate regions (pink, orange) with those from either end of red panda range extremes (red, yellow). Singletons were not included due to space limitations.

| Parsimony-informative position (35/340) | 2 | 5 | 10 | 29 | 36 | 52 | 55 | 80 | 81 | 82 | 83 | 86 | 87 | 90 | 92 | 94 | 99 | 100 | 104 | 106 | 139 | 150 | 154 | 161 | 162 | 196 | 213 | 221 | 245 | 268 | 281 | 286 | 291 | 293 | 300 |
|---|---|---|---|---|---|---|---|---|---|---|---|---|---|---|---|---|---|---|---|---|---|---|---|---|---|---|---|---|---|---|---|---|---|---|---|
| Consensus: | **T** | **T** | **C** | **C** | **C** | **C** | **G** | **A** | **T** | **T** | **A** | **A** | **G** | **A** | **T** | **T** | **G** | **C** | **T** | **C** | **T** | **G** | **C** | **T** | **A** | **C** | **C** | **C** | **C** | **C** | **G** | **T** | **A** | **A** | **C** |
| Haplotype name: | | | | | | | | | | | | | | | | | | | | | | | | | | | | | | | | | | | |
| SSP1 | | | | | | | | | | | | | | | C | | | | | | | A | | C | | | | | T | T | | | | G | |
| Dal3 | | | | | | | | | | | | | | | C | | | | | | | | T | | | T | | | T | | | | | G | |
| SSP3, Dal5 | | | | | | | | | | | | | | | C | | | | | | | A | | | | | | | T | T | | | | G | |
| SSP8, Li29, Hu28 | | | | | | T | | | | | | | | | C | | | | | | | | T | | | | | | T | | | | | G | |
| WRP-A, Dal1 | | | | | | | | | | | | | | | C | | | | | T | | | | | | | | | T | T | | | | G | |
| WRP-B, Dal2 | | | | | | | | | | | | | | | C | | | | | | | | T | | | | | | T | T | | | G | G | |
| WRP-D | | | | | | | | | | | | | | | C | | | | | | | | T | | | | | | T | | | | | G | |
| WRP-E | | | | | | | | | | | | | | | C | | | | | | | | T | | | | | | T | | | | | G | |
| WRP-F | | | | | | | | | | | | | | | C | | | | | | | | T | | | | | | T | | | | | G | T |
| WRP-G | | | | | | | | | | | | | | | C | | | | | | | | T | | | | | | T | | | | | G | |
| WRP-H | | | | | | | | | | | | | | | C | | | | | | | | T | | | | | | T | | | | | G | T |
| WRP-J | | | | | | | | | | | | | | | C | | | | | | | A | | C | | | | T | T | T | | | | G | |
| WRP-N | | | | | | | | | | | | | | | C | | | | | T | | | | | | | | | T | T | | | | G | |
| WRP-P | | | | | | T | | | | | | | | | C | | | | | | | | T | | | | | | T | | | | | G | |
| WRP-Q | | | | | | T | | | | | | | | | C | | | | | | | | T | | | | | | T | | | | | G | |
| WRP-R | | | | | | T | | | | | | | | | C | | | | | | | | T | | | | | | T | | | | | G | |
| WRP-S | | | | | | T | | | | | | | | | C | | | | | | | | T | | | | | | T | | | | | G | |
| SSP2, Aff-ref | | | | | | | | | | | | | | | C | | | | | | | | | | | | | | T | T | | | | G | |
| Dal7 | | | | | | T | | | | | | | A | | | | | | | | | | | | | | | | T | | | | | G | |
| Dal11 | | | | | | T | | | | | | | A | | | | | | | | | | | | | | | | T | | | | | G | |
| Dal12 | | | | | | T | | | | | G | | A | | | | | | | | | | | | | | | | T | | | | | G | |
| Dal13 | G | | G | | | T | | | | | | | A | | | | | | | | | | | | | | | | T | | | | | G | |
| Dal14 | G | | A | | T | G | | | | | | | A | | | | | | | | | | | | | | | | T | | | | | G | |
| Dal16 | | | | T | | T | | | | | | | A | | | | | | | | | | | | | | | | T | | | | | G | |
| Dal17 | | | | | | | | | | | | | A | | | | | | | | | | | | | | | | T | | | | | G | |
| Dal4 | | | | | | | | | | | | | | | | | | T | C | | C | A | | | | | | | T | | | | | G | |
| Dal6 | | | | | | | | | | | | | | | | | | T | C | | C | A | | | | | | | | | | | | | |
| WRP-C, Dal8 | | | | | T | | | | | | | | | | | | | T | C | | C | A | | | | | | | | | | | | | |
| Dal9 | | | | | T | | | | | C | | | | | | | | T | | | | A | | | G | | | T | | | | | | | |

| | | | | | | | | | | | | | | | | | | | | | | | | | | | | | | | | | | |
|---|---|---|---|---|---|---|---|---|---|---|---|---|---|---|---|---|---|---|---|---|---|---|---|---|---|---|---|---|---|---|---|---|---|---|
| Dal10 | | | | | | | | | C | | | | | G | | | | T | | | | | | | | T | T | | | | | | | |
| Dal15 | | | G | | | | | | C | | | | | | | | | T | | | | | | | | T | T | | | | | | | |
| Dal18 | | | | | | | | | | C | | | A | G | | | | T | | | | | | | | | | | | | | | | G |
| Li9 | | | | | | | | | | C | | | | G | | | | | | | | | | C | G | | | | | | | | | |
| Li11 | | | | | | | | | | C | | | A | G | | | A | | | | | | | | | | | | | T | | | | G |
| Li12 | | | | | | | | | | C | | | A | G | | | | T | | | | | | | | | | | | | | | | G |
| Li14, Hu5 | | | | | | | | | | C | | | | G | C | | | T | | | | | | | | T | | | | | | | | |
| Li16 | | | | | | | | | C | C | | | A | | | C | | | | | | | T | C | | | | | | | | | | |
| Li17 | | | | | | | | G | | | | G | | G | | | | T | | | | | | | G | ? | | | | | | | | |
| Li27 | | | | | | | | | | C | | | A | G | | | A | T | | | | | | | | | | | | T | | | | G |
| Li28, Hu1 | | | | | T | | | | | | | G | | G | | | | T | | | | | | | G | | | | | | | | | |
| Li 30 | | | | | | | A | | C | C | | | A | | | C | A | T | | | | | T | C | | ? | ? | | | | | C | | |
| Li31, Hu29 | | | | | | | | | | C | | | A | G | | | | | | | | | | | | | | | | | | | | G |
| Li32 | | | | | T | | | | | C | | G | | G | | | | T | | | | | | | G | | | | | | | | | |
| Li33 | | | | | T | | | | C | C | | | A | | | C | | | | | | | T | C | | | | | | | | | | |
| Li34 | | | | | T | | | | | C | | | A | G | | | | T | C | | | | | | | | | | | | | | | |
| Li35 | | C | | | | | | | C | | G | | | | | | | | | | | | | | | T | T | | | | | | | |
| Li36 | | | | T | | | | | | C | | | A | G | | | A | T | | | | | | | | | | | | | | | | G |
| Li 37, Hu25 | | | | | T | | | | | C | | | A | G | | | | T | | | | | | | | | | | | | | | | |
| Li38 | | | | | | | | | | C | | | | | | | | | | | | | | C | G | | | | | | | | | |
| Li39 | | | | | | | | | | C | | G | A | G | | | | T | | | | | | | | T | T | | | | | | | |
| Hu3 | | | | | | | | | C | C | | | A | | | C | | T | | | | | T | C | | | | | | | A | C | | G |
| Hu7 | | | | | T | | | G | | C | | | | | | | | T | | | | | T | | | A | | T | | | | | | |
| Hu8 | | | | | | | | | C | C | | | A | | | C | | | | | | | T | C | | | | | | | | | | |
| Hu10 | | | | | | | | | | C | | G | | G | | | | T | | | | | | | G | | | | | | | | | |
| Hu12 | | | | | T | | | | | | | | | | | | | T | | | | A | | | G | | | T | | | | | | |
| Hu13 | | C | | | | | | | C | | | | | G | | | A | | | | | | | | | T | T | | | | | | | |
| Hu14 | | | | | T | | | | | C | | | A | G | | | | T | C | | | | | | | | | | | | | | | |
| Hu15 | | | | | | | | | | | | | | | C | | | T | C | | | | | | | T | | | | | | | | |
| Hu16 | | | | | | | | | | C | | G | A | G | | | | T | | | | | | | | | | T | | | | | | |
| Hu17 | | | | | | | | | | C | | | | G | | | | | | | | | | C | G | | | | | | | | | |
| Hu18 | | | | | | | | | | C | | | A | G | | | A | T | | | | | | | | | | | | | | | | G |
| Hu19 | | | | | | | | | | C | | | A | G | | | A | | | | | | | | | | | | | T | | | | G |
| Hu21 | | | | | | | | | | C | | | A | G | | | | T | | | | | | | | | | | | | | | | G |
| Hu22 | | | | | | | | | C | | G | | | | | | | | | | | | | | | T | T | | | | | | | |
| Hu23 | | | | | | | A | | C | C | | | A | | | C | A | T | | | | | T | C | | | | | | | | C | | |
| Hu24 | | | | | | | | | C | | | | | G | | | | | | | | | | | | T | T | | | | A | | | |
| Hu26 | | | | | T | | | | | C | | | A | G | | | | T | C | | | | | | | | | | | | | | | |
| Hu27 | | | | | | | | | C | | G | | | | | | | T | | | | | | | | T | T | | | | | | | |
| SSP4, Li01, Hu11 | | C | | | | | | | C | | | | | G | | | | | | | | | | | | T | T | | | | | | | |

| | | | | | | | | | | | | | | | | | | | | | | | | | | | | | | | | | |
|---|---|---|---|---|---|---|---|---|---|---|---|---|---|---|---|---|---|---|---|---|---|---|---|---|---|---|---|---|---|---|---|---|---|
| SSP5, Hu4 | | | | | T | | | G | | | | G | | G | | | | T | | | | | | | G | | | | | | | | |
| SSP6, Li05, Li26, Hu6 | | | | | T | | | | | C | | G | A | G | | | | T | | | | | | | | | | T | | | | | |
| SSP9, Li13, Hu2 | | | | | T | | | | | C | | | | | | | A | T | | | | | | | | T | | | | | | | |
| SSP10 | | | | | T | | | G | | C | | | | | | | | T | | | | | T | | | A | | T | | | | | |
| SSP11, Hu20 | | | | | | | | | | | | G | | G | | | | T | | | | | | | G | | | | | | | | G |
| SSP7, Afs-ref, Hu9 | | C | | | | | | | C | | | | | G | | | A | | | | | | | | | T | T | | | | | | |

**Table C.** Number of samples used/study to determine each of 71 mtDNA haplotypes for population genetic analyses of native red pandas.

| H# | haplotypes: | # individuals/study: wild SSP^ | WRP | Li^^ | Hu | Dalui | **Combined** | (Li)* =Su |
|---|---|---|---|---|---|---|---|---|
| | *A. f. fulgens* | | | | | | | |
| 1 | SSP-1 | 12 | 0 | 0 | 0 | 0 | **12** | |
| 2 | SSP-3, Dal-5 | 5 | 0 | 0 | 0 | 3 | **8** | |
| 3 | SSP-8, Li-29, Hu-28 | 24 | 0 | 1 | 1 | 0 | **26** | |
| 4 | WRP-A, Dal-1 | 0 | 2 | 0 | 0 | 5 | **7** | |
| 5 | WRP-B, Dal-2 | 0 | 3 | 0 | 0 | 5 | **8** | |
| 6 | WRP-D | 0 | 9 | 0 | 0 | 0 | **9** | |
| 7 | WRP-E | 0 | 1 | 0 | 0 | 0 | **1** | |
| 8 | WRP-F | 0 | 1 | 0 | 0 | 0 | **1** | |
| 9 | WRP-G | 0 | 1 | 0 | 0 | 0 | **1** | |
| 10 | WRP-H | 0 | 2 | 0 | 0 | 0 | **2** | |
| 11 | WRP-J | 0 | 1 | 0 | 0 | 0 | **1** | |
| 12 | WRP-N | 0 | 1 | 0 | 0 | 0 | **1** | |
| 13 | WRP-P | 0 | 1 | 0 | 0 | 0 | **1** | |
| 14 | WRP-Q | 0 | 1 | 0 | 0 | 0 | **1** | |
| 15 | WRP-R | 0 | 1 | 0 | 0 | 0 | **1** | |
| 16 | WRP-S | 0 | 1 | 0 | 0 | 0 | **1** | |
| 17 | Dal-3 | 0 | 0 | 0 | 0 | 6 | **6** | |
| | **Subtotal:** | | | | | | **87** | |
| 18 | Dal-7 | 0 | 0 | 0 | 0 | 8 | **8** | |
| 19 | Dal-11 | 0 | 0 | 0 | 0 | 1 | **1** | |
| 20 | Dal-12 | 0 | 0 | 0 | 0 | 1 | **1** | |
| 21 | Dal-13 | 0 | 0 | 0 | 0 | 1 | **1** | |
| 22 | Dal-14 | 0 | 0 | 0 | 0 | 1 | **1** | |
| 23 | Dal-16 | 0 | 0 | 0 | 0 | 1 | **1** | |
| 24 | Dal-17 | 0 | 0 | 0 | 0 | 1 | **1** | |
| | **Subtotal:** | | | | | | **14** | |
| | ***A. f. f.* Total:** | - | - | - | - | - | **101** | |
| | *A. f. styani* | | | | | | | |
| 25 | Dal-4 | 0 | 0 | 0 | 0 | 1 | **1** | |
| 26 | Dal-6 | 0 | 0 | 0 | 0 | 1 | **1** | |
| 27 | WRP-C, Dal-8 | 0 | 2 | 0 | 0 | 3 | **5** | |
| | **Subtotal:** | | | | | | **7** | |
| 28 | SSP-4, Li-01, Hu-11 | 0 | 0 | 11 | 20 | 0 | **31** | 3 |
| 29 | SSP-5, Hu-4 | 0 | 0 | 0 | 9 | 0 | **9** | |
| 30 | SSP-6, Li-05, Li-26, Hu-6 | 0 | 0 | 7 | 20 | 0 | **27** | 4 |
| 31 | SSP-7, Hu-9, Yon-*Afs* ref | 0 | 0 | 0 | 10 | 0 | **10** | |
| 32 | SSP-9, Li-13, Hu-2 | 0 | 0 | 1 | 7 | 0 | **8** | |
| 33 | SSP-11, Hu-20 | 0 | 0 | 0 | 2 | 0 | **2** | |
| 34 | Li-09, Hu-17[ǂ] | 0 | 0 | 2 | 3 | 0 | **5** | 3 |
| 35 | Li-11, Hu-19[ǂ] | 0 | 0 | 1 | 1 | 0 | **2** | 1 |

| | | | | | | | | |
|---|---|---|---|---|---|---|---|---|
| 36 | Li-12 | 0 | 0 | 1 | 0 | 0 | **1** | 1 |
| 37 | Li-14, Hu-5 | 0 | 0 | 1 | 4 | 0 | **5** | 1 |
| 38 | Li-16** | 0 | 0 | 1 | 0 | 0 | **1** | 1 |
| 39 | Li-17 | 0 | 0 | 1 | 0 | 0 | **1** | 1 |
| 40 | Li-27 | 0 | 0 | 1 | 0 | 0 | **1** | |
| 41 | Li-28, Hu-1 | 0 | 0 | 2 | 6 | 0 | **8** | |
| 42 | Li-30 | 0 | 0 | 1 | 0 | 0 | **1** | |
| 43 | Li-31, Hu-29 | 0 | 0 | 1 | 2 | 0 | **3** | |
| 44 | Li-32 | 0 | 0 | 1 | 0 | 0 | **1** | |
| 45 | Li-33 | 0 | 0 | 1 | 0 | 0 | **1** | |
| 46 | Li-34 | 0 | 0 | 1 | 0 | 0 | **1** | |
| 47 | Li-35 | 0 | 0 | 1 | 0 | 0 | **1** | |
| 48 | Li-36 | 0 | 0 | 1 | 0 | 0 | **1** | |
| 49 | Li-37, Hu-25 | 0 | 0 | 1 | 1 | 0 | **2** | |
| 50 | Li-38 | 0 | 0 | 1 | 0 | 0 | **1** | |
| 51 | Li-39 | 0 | 0 | 1 | 0 | 0 | **1** | |
| 52 | Hu-3 | 0 | 0 | 0 | 2 | 0 | **2** | |
| 53 | Hu-7 | 0 | 0 | 0 | 3 | 0 | **3** | |
| 54 | Hu-8 | 0 | 0 | 0 | 3 | 0 | **3** | |
| 55 | Hu-10 | 0 | 0 | 0 | 5 | 0 | **5** | |
| 56 | Hu-12 | 0 | 0 | 0 | 2 | 0 | **2** | |
| 57 | Hu-13 | 0 | 0 | 0 | 1 | 0 | **1** | |
| 58 | Hu-14 | 0 | 0 | 0 | 1 | 0 | **1** | |
| 59 | Hu-15 | 0 | 0 | 0 | 1 | 0 | **1** | |
| 60 | Hu-16 | 0 | 0 | 0 | 3 | 0 | **3** | |
| 61 | Hu-18 | 0 | 0 | 0 | 3 | 0 | **3** | |
| 62 | Hu-21 | 0 | 0 | 0 | 3 | 0 | **3** | |
| 63 | Hu-22 | 0 | 0 | 0 | 1 | 0 | **1** | |
| 64 | Hu-23 | 0 | 0 | 0 | 1 | 0 | **1** | |
| 65 | Hu-24 | 0 | 0 | 0 | 2 | 0 | **2** | |
| 66 | Hu-26 | 0 | 0 | 0 | 1 | 0 | **1** | |
| 67 | Hu-27 | 0 | 0 | 0 | 1 | 0 | **1** | |
| 68 | Dal-9 | 0 | 0 | 0 | 0 | 2 | **2** | |
| 69 | Dal-10 | 0 | 0 | 0 | 0 | 2 | **2** | |
| 70 | Dal-15 | 0 | 0 | 0 | 0 | 1 | **1** | |
| 71 | Dal-18 | 0 | 0 | 0 | 0 | 1 | **1** | |
| | **Subtotal:** | - | - | - | - | - | **163** | |
| | ***A. f. s.* Subtotal:** | - | - | - | - | - | **170** | |
| | **TOTAL:** | **41** | **27** | **40** | **119** | **44** | **271** | **15** |

^ = previously determined haplotypes from zoo animals that were also found in wild Nepal population.

^^ = Several entries in Li's Table 2 are suspect; total # of individuals sampled differs because #40 not used.

* = + 22 Su samples with known origin: haplotype #2 (1), #3 (1), #4 (1), #6 (1), #7 (1), #8 (5), #10 (5), #15 (1), #18 (1), #19 (1), #20 (1), #21 (1), #24 (1), #25 (1) = 37; none used in this analysis except **.

**= used Su's haplotype? ǂ = newly found duplicate haplotype pair

**Table D.** Pairwise differences between two subspecies and four haplogroups of native red pandas (*N* = 271, *h* = 71), based on 340bp of mtDNA-CR. $F_{ST}$ is above diagonal and $\chi^2$ *P*-values below diagonal. Calculated individually in DnaSP6 (Gene Flow and Genetic Differentiation module) with option to exclude gaps only in pairwise comparisons and 1000 permutation test replicates.

| | ***A. f. fulgens*** | *Aff*-red | *Aff*-pink | ***A. f. styani*** | *Afs*-orange | *Afs*-yellow |
|---|---|---|---|---|---|---|
| *n* | 101 | 87 | 14 | 170 | 7 | 163 |
| *h* | 24 | 17 | 7 | 47 | 3 | 44 |
| ***A. f. fulgens*** | --- | 0.0095 | 0.4747 | **0.5224** | 0.6493 | 0.4777 |
| *Aff*-red | *0.9323ns* | --- | 0.3204 | 0.5609 | 0.6410 | 0.5215 |
| *Aff*-pink | *0.0007**** | *0.0000**** | --- | 0.5595 | 0.6590 | 0.5127 |
| ***A. f. styani*** | ***0.0000***** | *0.0000**** | *0.0000**** | --- | 0.5490 | -0.0047 |
| *Afs*-orange | *0.0000**** | *0.0000**** | *0.0090*** | *0.0000**** | --- | 0.5165 |
| *Afs*-yellow | *0.0000**** | *0.0000**** | *0.0000**** | *1.0000ns* | *0.0000**** | --- |

Astericks indicate significance level (*** = *P*<0.001, ** = 0.001<*P*<0.01, *ns* = not significant); *N/n* = # sequences, *h* = # haplotypes. Shaded values are not significant. **Bolded** values indicate comparison between subspecies.

**Table E.** Tests of neutrality of mutations for all native red panda groupings as defined by geographic locations and/or phylogeny, calculated in DnaSP6.

| Population (*h*) | Tajima's *D* *P* based on (*S*) | | Tajima's *D* *P* based on (*Eta*) | | Fu's $F_S$ ($F_S \leq$ obs.) | *P* (*h*=*h*) |
|---|---|---|---|---|---|---|
| ***A.f. fulgens*** (24) | -1.2059 (28) | *ns* | -1.3365 (30) | *ns* | -8.129 | 0.000 |
| *Aff* – red (17) | -0.7522 (18) | *ns* | -0.7522 (18) | *ns* | -4.798 | 0.005 |
| *Aff* – pink (7) | -1.5726 (9) | *ns* | **-2.0381 (11)** | ***** | -1.456 | **0.123** |
| ***A.f. styani*** (41)^^ | -0.0722 (35) | *ns* | -0.0722 (35) | *ns* | -13.597 | 0.000 |
| *Afs* – orange^ (3) | -0.3187 (4) | *ns* | -0.3187 (4) | *ns* | 0.789 | **0.346** |
| *Afs* – yellow (38)^^ | -0.0092 (33) | *ns* | -0.0092 (33) | *ns* | -11.489 | 0.000 |
| COMBINED (61)^^ | -0.3102 (50) | *ns* | -0.4111 (52) | *ns* | -25.339 | 0.000 |

Significance: *ns* = *P* > 0.10 or 0.10 > *P* > 0.05, * = P < 0.05, ** = *P* < 0.02. *** = P< 0.000;

*S* = segretating sites, *Eta* = total # of mutations, *h* = # of haplotypes, ^ = bare minimum # of sequences,

^^ = some haplotypes collapsed due to missing data in variable positions.

**Table F.** Analysis of molecular variance (AMOVA) to partition genetic variation of all studied native red pandas: **a)** between subspecies as determined by ML analysis in this study, and **b)** among and within four phylogeographic haplogroups, as calculated in Arlequin v. 3.5.2.2.

**a)**

| | d.f. | Observed Partition: variance components | Observed Partition: % of total variation | *P* | $F_{ST}$ |
|---|---|---|---|---|---|
| Between subspecies: | 1 | 2.577 Va | 48.0 | 0.0000 | **0.480** |
| Within subspecies: | 269 | 2.800 Vb | 52.0 | - - -- - - | |

d.f. = degrees of freedom; $F_{ST}$ = Fixation index; *P* = probability of having a more extreme variance component and $F_{ST}$ than the observed values by chance alone, based on comparisons with 1023 permutations per run.

**b)**

| | d.f. | Observed Partition: variance components | Observed Partition: % of total variation | *P* | $F_{ST}$ |
|---|---|---|---|---|---|
| Among haplogroups: | 3 | 2.749 Va | 52.0 | 0.0000 | **0.520** |
| Within haplogroups: | 267 | 2.539 Vb | 48.0 | - - - - - - | |

d.f. = degrees of freedom; $F_{ST}$ = Fixation index; *P* = probability of having a more extreme variance component and $F_{ST}$ than the observed values by chance alone, based on comparisons with 1023 permutations per run.

#### SUPPLEMENTARY INFORMATION

##### II. Preliminary results from phylogenetic analyses of individual studies to determine duplicate haplotypes

Missing links connect the phylogeographic structure of endangered red pandas,
remaining as one species – *Ailurus fulgens*, and expediting conservation

**Biodiversity and Conservation**

Lucy A. Dueck[1] and Deniz Aygören Uluer

Su et al.'s (2001) haplotypes all sorted out as the *A. f. styani* subspecies, except possibly one – Su21 from sample Rn, in Gongshan, northernmost Yunnan near Myanmar (two other samples – Su-17, Su-20) from the same location were identified as *A. f. styani*, though). Upon visual inspection, this haplotype shared bases at variable positions with both subspecies (but more with *A. f. fulgens*), which could place Su21 between subspecies on a haplotype network. In the unrooted NJ phylogram (**Figure A**), Su21 identified as an ancestor to all *A. f. fulgens* sequences in this comparison, but this finding must be regarded cautiously due to poor support and short segments. A discrepancy about the affinity of a subclade containing putative *A. f. styani* haplotypes SSP-4 and 7, Su-01, -07, -08, and -23, and reference sample Yonezawa was evident, but likely due to false signal from a very abbreviated sequence segment of 219 bp analyzed.

Each of the two subsequent studies by Chinese authors yielded only one haplotype that was likely *A. f. fulgens* – Li et al.'s (2005) Li-29, one of two samples from Zhangmu, East? Tibet, and Hu et al.'s (2011) Hu-R28, which matched Li-29 (neither phylogram shown, but see below). Hu-R28 was obtained from museum skin traced back to either of two areas in southeast Tibet/ Arunachal Pradesh? just east of the Siang River. The other two samples collected from the same areas (Li-31 and Hu-R29) sorted out as *A. f. styani*, however.

In our individual analysis of Dalui et al.'s (2021) haplotypes, three (H4, H6, H8) aligned with *A. f. styani* instead of with their *A. f. fulgens* subclade 1c (H8 matching WRP-C). However, four other haplotypes from the same region concurred in alignment with their *A. f. fulgens*. All seven Indian haplotypes were found in Sikkim and West Bengal between Nepal and Bhutan, near where both subspecies were also found in eastern Nepal by Dueck and Steffens (2022).

To further investigate this major switch of affinities for Dalui et al.'s (2021) subclade 1c, the three Chinese sequences that proved to be *A. f. fulgens* in our individual analyses were added to the dataset and results shown as an unrooted NJ phylogram (**Figure Ba**). Both Chinese haplotypes from Tibet (Hu-R28, Li-29), used by Dalui et al. (2021) for their haplotype H19 in subclade 1b again aligned with *A. f. fulgens* here. Su21 from Yunnan aligned with Dalui's *A. f. fulgens* subclade 1a and matched Dalui-H11 in the 175 sequences that overlapped. This Dalui subclade 1a (pink box) was found east of Bhutan, aligning with *A. f. fulgens* in the western part of the range. But the Dalui subclade 1c (orange box), containing WRP-C and Dalui H4-6-8, was found west of Bhutan, aligning with *A. f. styani* in the eastern part of the species' range.

More analyses were then undertaken to place these two subclades relative to the red panda subspecies. Since Dueck (2021) found the most reliable phylogeny was produced using 10 outgroups due to long-branch attraction issues, a phylogram in radial form so rooted is presented here as **Figure Bb**. It demonstrates that these two subclades are distinct but perhaps intermediary between the two more tightly related groups of subspecies.

**Fig. A** Unrooted, uncondensed NJ phylogram of Su et al.'s (2001) 23 mtDNA haplotypes (excluding 2 with many missing bases) were collected from zoo/museum specimens with known origins in two Chinese provinces, combined with 11 SSP (US zoos) and 14 WRP (Nepal) haplotypes, plus two reference sequences (one for each subspecies). The blue dashed line indicates the division between subspecies as determined by this analysis. The haplotype of interest (Su21) is enclosed by an oval. This bootstrap consensus tree, inferred from 10,000 replicates, represents the evolutionary history of the taxa analyzed using the Neighbor-Joining method. The tree is drawn to scale, the evolutionary distances were computed using the Tamura-Nei method and in units of the number of base substitutions per site. The percentages of replicate trees in which the associated taxa clustered together are shown on the branches. Codon positions with alignment gaps and missing data were eliminated only in pairwise sequence comparisons, with a total of 219 positions used (two sequences that were missing many bases in the front [Su16, Su19] were eliminated). Phylogenetic analysis was conducted in MEGA4.

**Fig. Ba** Unrooted NJ phylogram of Dalui et al.'s (2021) 18 mtDNA haplotypes collected in the wild from three states in northeastern India, combined with 11 SSP (US zoos) and 14 WRP (Nepal) haplotypes, two reference sequences (one for each subspecies), and 3 sequences-of-note from each of three previous Chinese studies on red panda genetic diversity. The blue dashed line indicates the division between subspecies as determined by this analysis. Each clade or subclade as defined by Dalui et al. (2021) is enclosed in a large box; those of phylogenetic interest are colored pink or orange. Haplotypes of interest are enclosed by ovals. This bootstrap consensus tree, inferred from 10,000 replicates, represents the evolutionary history of the taxa analyzed using the Neighbor-Joining method. The tree is drawn to scale, and the evolutionary distances were computed using the Tamura-Nei method and in units of the number of base substitutions per site. Codon positions with alignment gaps and missing data were eliminated only in pairwise sequence comparisons, with a total of 339 positions used. Phylogenetic analyses were conducted in MEGA4. **Fig. Bb** Radial phylogram of the same 48 haplotypes as above, but rooted by the 10 outgroups used in Dueck (2021) to balance long-branch attraction. This phylogram also has the two Dalui subclades of interest demarcated by: pink triangle for subclade 1a/Su21 and by orange oval for subclade 1c/WRP-C. This analysis was computed exactly as above except that it was based on 348 positions used.

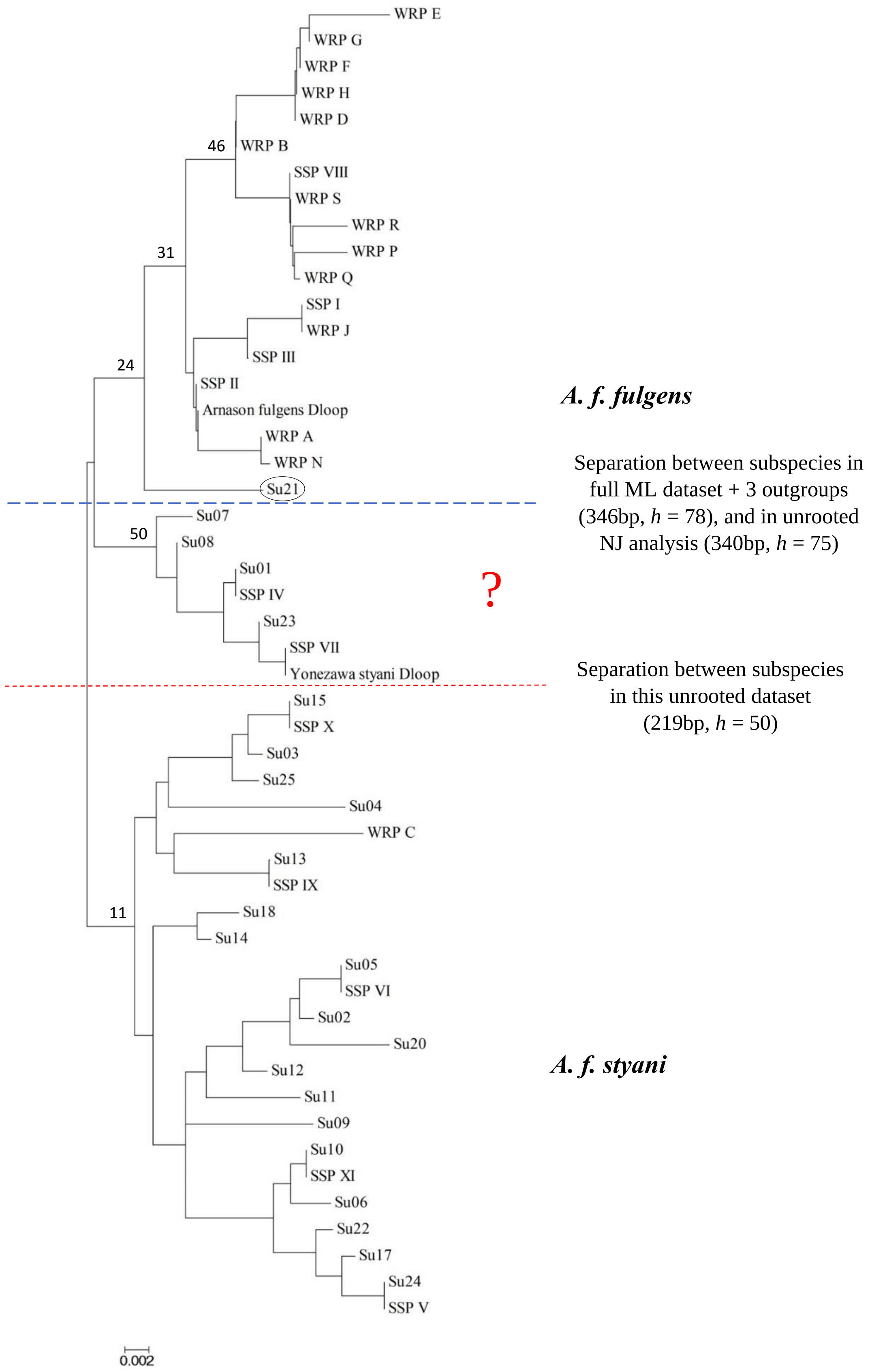

WRP E
WRP G
WRP F
WRP H
WRP D
46
WRP B
SSP VIII
WRP S
WRP R
WRP P
31
WRP Q
SSP I
WRP J
SSP III
24
SSP II
Arnason fulgens Dloop
WRP A
WRP N
Su21
Su07
50
Su08
Su01
SSP IV
Su23
SSP VII
Yonezawa styani Dloop
Su15
SSP X
Su03
Su25
Su04
WRP C
Su13
SSP IX
11
Su18
Su14
Su05
SSP VI
Su02
Su20
Su12
Su11
Su09
Su10
SSP XI
Su06
Su22
Su17
Su24
SSP V
0.002
A. f. fulgens
Separation between subspecies in full ML dataset + 3 outgroups (346bp, h = 78), and in unrooted NJ analysis (340bp, h = 75)
?
Separation between subspecies in this unrooted dataset (219bp, h = 50)
A. f. styani

**Fig. A**

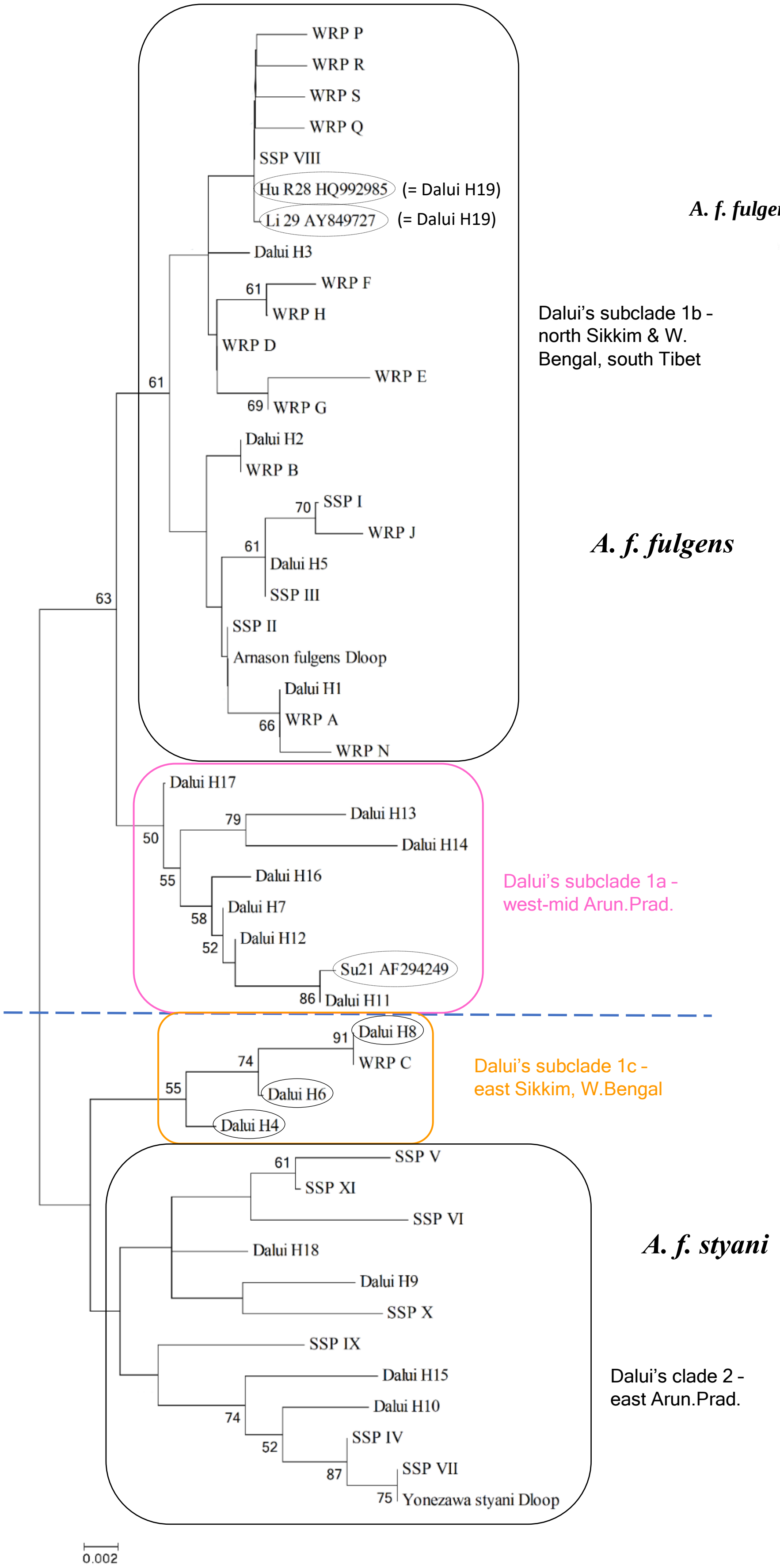
WRP P
WRP R
WRP S
WRP Q
SSP VIII
Hu R28 HQ992985
(= Dalui H19)
Li 29 AY849727
(= Dalui H19)
Dalui H3
WRP F
WRP H
WRP D
WRP E
WRP G
Dalui H2
WRP B
SSP I
WRP J
Dalui H5
SSP III
SSP II
Arnason fulgens Dloop
Dalui H1
WRP A
WRP N
Dalui's subclade 1b -
north Sikkim & W.
Bengal, south Tibet
A. f. fulgens
Dalui H17
Dalui H13
Dalui H14
Dalui H16
Dalui H7
Dalui H12
Su21 AF294249
Dalui H11
Dalui's subclade 1a -
west-mid Arun.Prad.
Dalui H8
WRP C
Dalui H6
Dalui H4
Dalui's subclade 1c -
east Sikkim, W.Bengal
SSP V
SSP XI
SSP VI
Dalui H18
Dalui H9
SSP X
SSP IX
Dalui H15
Dalui H10
SSP IV
SSP VII
Yonezawa styani Dloop
A. f. styani
Dalui's clade 2 -
east Arun.Prad.
0.002

Fig. Ba

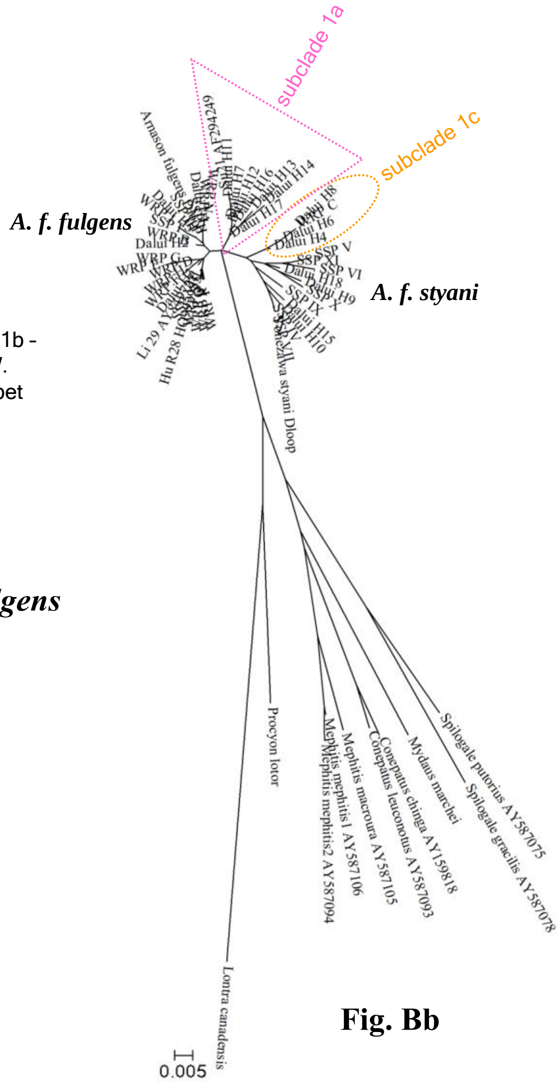
subclade 1a
subclade 1c
A. f. fulgens
A. f. styani
Yonezawa styani Dloop
Procyon lotor
Mephitis mephitis1 AY587106
Mephitis mephitis2 AY587094
Mephitis macroura AY587105
Conepatus leuconotus AY587093
Conepatus chinga AY159818
Mydaus marchei
Spilogale putorius AY587075
Spilogale gracilis AY587078
Lontra canadensis
0.005

Fig. Bb

**SUPPLEMENTARY INFORMATION**

#### III. Why are there still midrange gaps in the genetic assessment of red pandas?

Missing links connect the phylogeographic structure of endangered red pandas, remaining as one species – *Ailurus fulgens*, and expediting conservation

**Biodiversity and Conservation**

Lucy A. Dueck[1] and Deniz Aygören Uluer

There have been several studies on suitable habitat, potential distribution, threats, and conservation of red pandas in Bhutan (Yonzon et al. 1997; Dorji et al. 2012; Wangchuk 2014; Tobgay and Mahavik 2020; Letro et al. 2022; Dendup et al. 2023; among others). The most recent two publications revealed that red panda habitat is present in 19 of 20 districts, 8 of 11 protected areas, and 5 of 8 biological corridors in the country (almost 30% of the total area). Bhutan is fortunate to not only be a biodiversity hotspot, but also in having a strong environmental ethic to preserve its natural resources in several large protected areas connected by corridors. They also have robust legislation via the Forest and Nature Conservation Act 1995 that strictly protects red pandas by listing them under Schedule I, as well as cultural implications against harming them. Conservation actions specific to red pandas have been limited however, only effected by broad landscape or umbrella species conservation, until the recent development of Bhutan's Red Panda Conservation Action Plan (2018-2023). Unfortunately, it makes no allowance for genetic assessment of red pandas in Bhutan, which has never been undertaken. The reasons for this deficit are complex – the government is very protective of its natural resources and committed to environmental conservation. Regulations restrict external researchers from conducting studies, especially genetic, unless they align with Bhutan's conservation priorities and involve collaboration with local entities. Yet Bhutan does not have the funding, facilities, or expertise to conduct genetic studies on their own (Sangay Dorji, PhD, formerly of the Nature Conservation Division, Bhutan, and author of several publications on ecology of red pandas in Bhutan; pers. comm.). However, a recent announcement was made that the use of eDNA for biodiversity monitoring is being developed since 2022 through stakeholders WWF Bhutan, Department of Forests and Park Services, College of Natural Resources at the Royal University of Bhutan, ETH Switzerland, and others (Nature Conservation Division, Facebook, October 27, 2023 post, accessed 13 Feb. 2024). Perhaps this technology can be further advanced to initiate genetic assessments on red pandas in Bhutan.

No genetic studies of red pandas exclusively from Myanmar (formerly Burma) have been published either. Only nine Burmese red pandas have been sampled in broader studies included in our dataset – seven from museum skins by Li et al. (2005), and two by Hu et al. (2011) but inseparable from their Yunnan Gaoligong population; all clustered with the eastern *Afs* lineage. The only suitable red panda habitat in Myanmar lies to the north at the eastern extent of the Himalayan foothills. Red panda presence has been confirmed in three protected areas, but principally within and surrounding Imawbum National Park. A focused study in this area bordering Yunnan was conducted from 2010 to 2018 to assess conservation status via interviews, transects, and camera trapping. They found that hunting for trade with China, as well as habitat degradation and increased access from road building for illegal logging since 2000, posed the biggest threats to red pandas in Myanmar. Interestingly, their photo of two red pandas together in the snow showed noticeable differences between them in pelage color of tail and forehead (Lin et al. 2022). Difficult access and political unrest likely contribute to the dearth of any research other than the above on red pandas in Myanmar.

- - - - - - - - - - - - - - - - - - - - - - - - - - - - - - - - - - - - - - - - - - - - -

### SUPPLEMENTARY INFORMATION

#### IV. Photographs

Missing links connect the phylogeographic structure of endangered red pandas, remaining as one species – *Ailurus fulgens*, and expediting conservation

**Biodiversity and Conservation**

Lucy A. Dueck[1] and Deniz Aygören Uluer

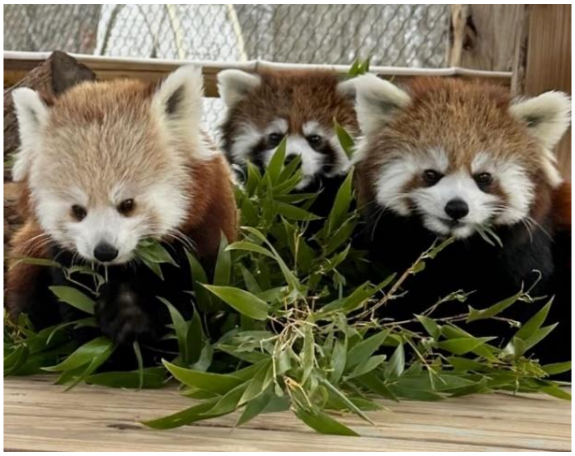

**A)** Photo by Keeper Kelly

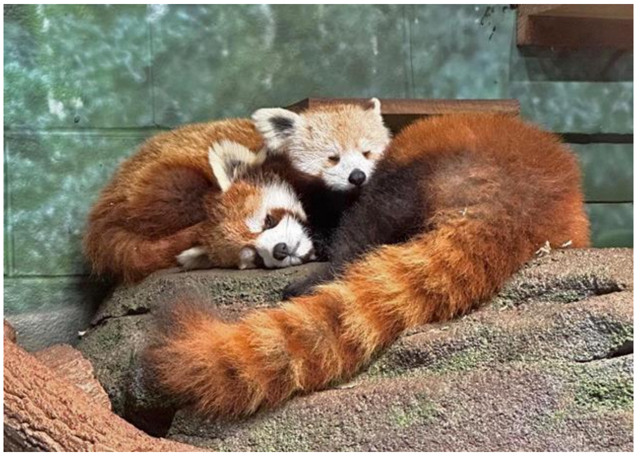

**B)** Photo by Keeper Dakota

**Photos A** (left) and **B** (right) of cubs from both red panda 'subspecies' being raised together at Greensboro Science Center, NC, posted on Facebook in Jan. 2024 and May 2024, respectively. The younger *A. f. fulgens* cub Miso (light forehead, both photos) was born at the Smithsonian National Zoo and transported to GSC for socialization with *A. f. styani* cubs Zuko (**A**) and Azula (**A**, **B**). Used with permission of photographers.
